# Cannabinoids shift the basal ganglia microRNA m^6^A methylation profile towards an anti-inflammatory phenotype in SIV-infected rhesus macaques

**DOI:** 10.1101/2024.10.11.614514

**Authors:** Chioma M. Okeoma, Lakmini S. Premadasa, Chen S. Tan, Ionita C. Ghiran, Mahesh Mohan

## Abstract

**Background:** Epitranscriptomic modifications modulate diverse biological processes like regulation of gene expression, abundance, location and function. N6-methyladenosine (m^6^A) methylation has been shown to regulate various diseases, including cancer and inflammation. While there is evidence that m^6^A modification is functionally relevant in neural development and differentiation, the role of m^6^A modification in HIV neuropathogenesis is unknown.

**Methods:** Here, we used anti-N6-methyladenosine (m^6^A) antibody immunoprecipitation and microarray profiling to identify m^6^A modifications in miRNAs in basal ganglia (BG) of Rhesus macaques (RMs) that were uninfected (VEH) and SIV-infected on combination anti-retroviral therapy (cART) and either VEH-treated (VEH/SIV/cART), or THC:CBD-treated (THC:CBD/SIV/cART). Ingenuity pathway analysis was conducted to understand the biological implications of miRNA m^6^A methylation in HIV neuropathogenesis. Finally, to understand the functional significance of m^6^A modifications in miRNAs, we overexpressed FAM-labeled wild-type or m^6^A-modified miR-194-5p in SCC-25 cells and determined its impact on the expression of its target, STAT1, an interferon-stimulated transcription factor known to drive persistent neuroinflammation in several neurodegenerative diseases.

**Results:** HIV/SIV infection promoted an overall hypomethylated miRNA m^6^A profile. While the overall hypomethylated m^6^A profile was not significantly impacted by THC:CBD, specific miRNAs predicted to target proinflammatory genes showed markedly reduced m^6^A methylation levels compared to the VEH-treated RMs. Additionally, specific BG tissue miRNAs bearing m^6^A epi-transcriptomic marks were transferred and detected in BG-derived extracellular vesicles. Mechanistically, the DRACH motif in the seed region of miR-194-5p was significantly m^6^A hypomethylated in THC:CBD/SIV/cART RMs. In SCC-25 cells, unlike wild-type miR-194-5p, transfected m^6^A-modified miR-194-5p mimics failed to downregulate STAT1 protein expression. Further, compared to VEH/SIV/cART RMs, THC:CBD administration significantly reduced m^6^A methylation of 44 miRNAs directly involved in regulating CNS network genes.

**Conclusions:** These results underscore the need for investigating the qualitative, and posttranscriptional modifications in RNA along with the more traditional, quantitative alterations in pathological conditions or in response to disease modifying treatments. Our findings indicate that m^6^A epitranscriptomic marks in the seed nucleotide region can impair miRNA function and that cannabinoids may preserve it by reducing m^6^A methylation levels. Finally, these findings provide a novel mechanistic (miRNA m^6^A hypomethylation) explanation underlying the anti-neuroinflammatory effects of phytocannabinoids in HIV/SIV infection.

## Introduction

Despite successful control of HIV with combined antiretroviral therapy (cART)[1–3], up to 50% of persons living with HIV (PLWH) still suffer from HIV Associated Neurocognitive Disorders (HAND)[4, 5]. A major factor driving HAND is incomplete viral suppression in the CNS, and the consequent activation of resident microglia, leading to neuroinflammation and neuronal damage.

In addition, drug abuse is a major comorbidity, where ∼40% of PLWH use cocaine and cannabis[6–9], resulting in increased virus-induced pathology[10–15], though cannabis use could afford varying degrees of protection from immune activation/inflammation [7]. While collecting longitudinal blood and matched brain samples from PLWH and healthy controls is extremely challenging, the availability of the SIV-infected RM model supports investigation of phytocannabinoids and other drugs of abuse on HIV/SIV neurological disease progression and has the potential to shed light on the underlying molecular mechanisms. Our previous studies, using the non-human primate (NHP) model, showed that twice daily administration of low-dose delta-9-tetrahydrocannabinol (THC) to RMs significantly reduced expression of type I-interferon (IFN) induced genes including inflammation associated miRNAs, miR-155 and miR-142-3p, in basal ganglia (BG) of cART naïve SIV-infected RMs[16, 17]. The effect of THC on BG was also present in BG-derived extracellular vesicles (EVs). Transcriptomic analyses of miRNAs associated with EVs isolated from these BG tissues revealed that both SIV infection and THC administration induced distinct but significant changes in BG EV-associated miRNAs, which were predicted to regulate pathways related to inflammation/immune regulation, TLR signaling, neurotrophin TRK receptor signaling, and cell death/response[18]. Long-term administration of low-dose THC to SIV infected RMs led to significant upregulation of 37 miRNAs in BG-EVs. Of these 37 miRNAs, 11 miRNAs were significantly downregulated in EVs from SIV infected control RMs that received vehicle control instead of THC.

Recent studies have identified over 170 RNA modifications, among which, m^6^A (N6-methyladenosine) is the most abundant modification in mammalian RNA, occurring in ∼50% of the total methylated ribonucleotides[19]. RNA m^6^A modifications regulate a wide variety of processes such as metabolism, splicing, translation, and degradation of RNA as well as in disease pathogenesis. m^6^A modifications have been described not only in mRNA but also recently in miRNAs, a class of small noncoding regulatory RNAs. miRNAs play key roles in regulating protein translation and has been implicated in various neurological disorders, including cognitive impairment. While others and we have published the role of miRNAs in the regulation of gene expression in the brain, there is a significant knowledge gap regarding the role of post transcriptional changes, in particular, miRNA m^6^A modification and its implications in HIV/SIV neuropathogenesis. Accordingly, the present study investigated the relationship between HIV/SIV infection of BG, miRNA m^6^A alterations, and the potential impact of concurrent phytocannabinoid use on miRNA m^6^A modifications and its potential implications for HAND pathogenesis.

Here, we present data supporting shifts in the small RNA landscape in the BG in response to HIV/SIV infection and cART treatment and the impact of concurrent THC:CBD administration. In addition to changes in the expression levels of several miRNA species involved in cognition and neuroinflammation, we also detected for the first-time significant alterations in the m^6^A methylation levels of key miRNA species, suggesting an added, more refined level of protein expression control. By exploring the interplay involving HIV/SIV infection, miRNA m^6^A alterations in BG, and phytocannabinoid use, this study aims to offer a comprehensive understanding of the role of m^6^A modifications on miRNA function that may provide valuable mechanistic insights into the development of HAND and pave the way for novel therapeutic strategies to mitigate cognitive decline in PLWH.

## Methods

### Animal care, ethics, and experimental procedures

All experiments using RM (**Table 1**) were approved by the Texas Biomedical Research Institute’s (Texas Biomed) Institutional Animal Care and Use Committee (Protocols 1734MM). The Southwest National Primate Research Center (SNPRC) is an association for Assessment and Accreditation of Laboratory Animal Care International-accredited facility (AAALAC #000246). The NIH Office of Laboratory Animal Welfare assurance number for the SNPRC is D16-00048. All clinical procedures, including administration of anesthesia and analgesics, were carried out under the direction of a laboratory animal veterinarian. Animals were pre-anesthetized with ketamine hydrochloride, acepromazine, and glycopyrrolate, intubated and maintained on a mixture of isoflurane and oxygen. All possible measures were taken to minimize the discomfort of all the animals used in this study. Texas Biomed complies with NIH policy on animal welfare, the Animal Welfare Act, and all other applicable federal, state and local laws.

**Table 1.** Animal IDs, SIV inoculum, duration of infection and viral loads in combination anti-retroviral therapy treated chronic SIV-infected rhesus macaques receiving vehicle or Δ^9^-THC :CBD combination.

| Table 1. Animal IDs, SIV inoculum, duration of infection and viral loads in combination anti-retroviral therapy treated chronic SIV-infected rhesus macaques receiving vehicle or $\Delta^9$-THC :CBD combination | | | | | | |
| --- | --- | --- | --- | --- | --- | --- |
| Animal ID | SIV Inoculum | Duration of Infection (days) | 12 DPI Plasma viral loads $10^6$/mL | Plasma Viral loads at necropsy $10^6$/mL | Basal Ganglia viral loads $10^6$/mg | Opportunistic Infections |
| Chronic SIV-Infected, cART and Vehicle treated |  |  |  |  |  |  |
| VSART1 | SIVmac251 | 180 | 2.50 | ND | ND | ND |
| VSART1 | SIVmac251 | 180 | 5.43 | ND | ND | ND |
| VSART1 | SIVmac251 | 180 | 1.93 | ND | ND | ND |
| VSART1 | SIVmac251 | 180 | 2.23 | ND | ND | ND |
| Chronic SIV-Infected, cART and $\Delta^9$-THC:CBD treated | | | | | | |
| TCART1 | SIVmac251 | 180 | 8.2 * | ND | ND | ND |
| TCART2 | SIVmac251 | 180 | 15.8 * | ND | ND | ND |
| TCART3 | SIVmac251 | 180 | 16.8 # | ND | ND | ND |
| Uninfected Controls |  |  |  |  |  |  |
| Control 1 | NA | NA | NA | NA | NA | NA |
| Control 2 | NA | NA | NA | NA | NA | NA |
| Control 3 | NA | NA | NA | NA | NA | NA |
| Control 4 | NA | NA | NA | NA | NA | NA |

ND- None detected
NA- Not application
\* - 20 DPI viral load

### Animal model and experimental design

Twelve age and weight-matched male Indian RMs were randomly distributed into three groups (**Table 1**). Group 1 [uninfected controls (n = 4)] served as uninfected controls. Groups 2-3 (n = 38) were infected intravenously with 100 times the 50% tissue culture infective dose (100TCID_50_) of SIVmac251. Group 2 [VEH/SIV/cART] (n = 4) received twice daily injections of vehicle (VEH) (1:1:18 of emulphor:ethanol:saline), intramuscularly. Group 3 [THC:CBD/SIV/cART] (n = 4) received twice daily injection of THC:CBD (1:3 ratio) intramuscularly beginning four weeks after SIV infection at 0.18mg/kg, as used in previous studies[17]. This dose of THC was increased to 0.32mg/kg over a period of 2 weeks and maintained for the duration of the study. For animals in groups 2 and 3 (n = 8), cART was administered daily by subcutaneous injection and included TDF (5.1 mg/kg) (Gilead), Emtricitabine (30mg/kg) (Gilead) and Dolutegravir (3mg/kg) (ViiV Healthcare) and was initiated four weeks post-SIV infection (same time as initiation of cART or VEH injections). Basal ganglia (BG) tissue (∼5cm) was collected at necropsy from all animals. A cube of 1x1x1 cm BG tissue was used for total RNA extraction. For histopathological and immunohistochemical evaluation, BG tissues were fixed in Z-fix, embedded in paraffin, sectioned at 5 µM for further analysis. SIV levels in basal ganglia were quantified using the TaqMan One-Step Real-time RT-qPCR assay that targeted the LTR gene[16]. SIV inoculum, infection duration, BG viral loads, and BG histopathology is provided in **Table 1**.

### Methylated RNA immunoprecipitation (MeRip)

We have completed the Arraystar Human N6-methyladenosine (m6A) small RNA modification microarray analysis of the samples you submitted. The RNA samples were QC’d for quantity by NanoDrop ND-1000 spectrophotometer and RNA integrity by Bioanalyzer 2100 or gel electrophoresis. Briefly, 1-5 μg of each total RNA sample was immunoprecipitated by 4 μg anti-N6-methyladenosine antibody (Synaptic Systems) with 1 mg Protein G Dynabeads (Thermo Fisher, 11203D) in 500 μL RIP buffer. The modified RNAs were eluted from the immunoprecipitated magnetic beads as the “IP”. The unmodified RNAs were recovered from the supernatant as “Sup”. The “IP” and “Sup” RNAs were enzymatically labeled with Cy5 and Cy3 respectively in separate reactions using Arraystar’s standard protocols. The labeled RNAs were combined and hybridized onto Arraystar Human small RNA Modification Microarray (8x15K,

Arraystar) and the array was scanned in two-color channels by an Agilent Scanner G2505C.Agilent Feature Extraction software (version 11.0.1.1) was used to analyze the acquired array images. Raw intensities of IP (immunoprecipitated, Cy5-labelled) and Sup (supernatant, Cy3-labelled) were normalized with average of log2-scaled Spike-in RNA intensities. After normalization, the probe signals having Present (P) or Marginal (M) QC flags in at least 4 out of 12 samples were retained. Multiple probes from the same small RNA (miRNA/TsRNA (tRF&tiRNA)/pre-miRNA) were combined into one RNA level. “m6A abundance” was analyzed based on the Cy5-labelled IP (modified RNA) normalized intensities. Differentially m6A-modified RNAs between two comparison groups were identified by fold change (FC) and statistical significance (p-value) thresholds. Hierarchical Clustering heatmap was plotted to display m6A-modification patterns among samples by R software.

### RNA processing and m^6^A analyses

Basal ganglia tissues were collected during necropsy in 1 cm^3^ cubes and processed for total RNA extraction using Qiazol (Qiagen Inc, CA). A detailed protocol is provided in the Supplemental methods. Briefly, total RNA was immunoprecipitated with anti-N6-methyladenosine (m6A) antibody and the modified RNAs were eluted from the beads as "m6A," and unmodified RNAs were recovered as "unmodified" samples. Raw intensities of each sample were normalized with average of log2-scaled Spike-in RNA intensities. Differentially m6A-modified RNAs between two comparison groups were identified by fold change (FC) and statistical significance (p-value) thresholds. Hierarchical Clustering heatmap was plotted to display m6A-modification patterns among samples by R software.

### Criteria for selecting m^6^A modified miRNA

Differentially significant m^6^A methylated miRNAs were identified with p-value < 0.05, | log2(FC)| > 1 upregulation, <-1 downregulation). To identify m^6^A modified miRNAs that are biologically relevant to HIV/SIV induced neuroinflammation, we used miRNA Target Filter functionality of IPA with a set of criteria to prioritize experimentally validated and predicted mRNA targets to perform enrichment analysis. Based on confidence level, diseases, cells/tissues, and pathways, we identified miRNA-mRNA, overlay miRNA data onto network pathways, and molecules. The identified miRNA target genes were fed into WEB-based GEne SeT AnaLysis Toolkit (WebGestalt, https://www.webgestalt.org/) and used for Geneontology analysis. The following criteria were used:

Tissue = Brain

Pathway = Neurotransmitters and other nervous system signaling

Disease = neurological diseases

Network overlay = neuroinflammation canonical signaling pathway. Merge multiple relevant networks.

Used IPA-generated list for STRING: functional protein association networks (https://string-db.org/) for protein:protein analysis.

Used IPA-generated list for WEB-based Gene SeT AnaLysis Toolkit (WebGestalt) GO biological, cellular, molecular processes analyses, using over representation analysis (ORA) enrichment method.

### Cell culture and in vitro transfection studies

We determined the impact of miR-194-5p m6A on STAT1 using human squamous cell carcinoma (SCC-25) epithelial-like cell line (ATCC.org CAT# CRL-1628) as this cell line expressed high basal levels of STAT1 protein. SCC-25 cells were cultured in 1:1 mixture of DMEM and F12 medium (Thermo Fisher scientific, Waltham, MA USA) containing 1.2g/L sodium bicarbonate, 2.5mM L-glutamine, 15 mM HEPES and 0.5 mM sodium pyruvate and supplemented with 10% FBS in 8 well chamber slides (Cellvis, Mountain view, CA USA) at 37 °C in a humidified atmosphere with 5% CO_2_. At 90% confluency, cells were transfected with 30 nM of negative control, miR-194-5p-Wild type or miR-194-5p-m6A mimic using the Lipojet transfection reagent (Signagen, DE). Each treatment was performed in five replicates and repeated twice. At 72 h post-transfection, cells were fixed with 2% paraformaldehyde and immunostained with STAT1 and later with DAPI for nuclear localization.

### Immunofluorescence staining, confocal microscopy and data analysis

Immunofluorescence staining for the detection of STAT1 in SCC-25 cells transfected with negative control, miR-194-5p wild type and miR-194-5p m6A modified mimics was performed using STAT1 monoclonal antibody (Cell Signaling technology; Cat# 14994T) (1:200 dilution). A total of 5 images were taken from each well using Zeiss LSM700 confocal microscope (Carl ZEISS Microscopy, LLC) at 20X objective. Digital images were imported into HALO software (Indica Labs) for image quantitation analysis. Area quantification module FL v2.3.4 on HALO v3.6 (Indica Labs) was used to quantify STAT1 fluorescence intensity (red signal/Alexa-568). Artificial intelligence driven HALO software identifies all cells that express STAT1 in red and nuclei in blue (DAPI) and categorizes the cells based on fluorescence intensity levels. The average fluorescence intensity per each transfected well (5 replicates per condition) in 8 well chambers was calculated, and the data were graphed using Prism v9 (GraphPad Prism software).

### Statistical analysis

Pie Charts were generated using GraphPad Prism. Heatmaps were generated with GraphPad Prism and Heatmapper (http://heatmapper.ca/). Venn diagrams were generated with Venny 2.1 (https://bioinfogp.cnb.csic.es/tools/venny/). Changes in STAT1 protein expression in response to miR-194-5p transfection was analyzed using One-way ANOVA. Post hoc analysis was performed using Tukey’s multiple comparison’s test. Shapiro-Wilk test (GraphPad Prism) was used to test for data normality.

## Results

### Quantification of m^6^A profiles of BG miRNAs point to mechanistically relevant species involved in the anti-inflammatory effect of THC:CBD treatment

Altered m6A-modified RNA expression levels in the brain were linked to dysregulation of key genes involved in immune cell activation, cytokine production, and blood-brain barrier integrity, which could modulate the activation of microglia and astrocytes, thereby, initiating and exacerbating neuroinflammatory processes[20, 21]. Most of the published research regarding the role of m6A changes in neuroinflammation has centered on mRNA and lncRNA and not miRNA molecules (reviewed in[22]. Hence, we focused exclusively on miRNAs in the present study.

All seven SIV-infected RMs, receiving either VEH (n=4) or THC:CBD (n=3), had substantial plasma viral loads ranging from 1.93 to 16.8 x 10^6^ copies/mL during acute infection [12-20 days post SIV infection (DPI)] (**Table 1**). However, cART initiation at 30 DPI effectively suppressed viral replication, as viral RNA was undetectable in both plasma and BG at necropsy (180 DPI).

We investigated how SIV infection and/or THC:CBD altered the landscape of m^6^A changes in miRNAs isolated from BG of RMs that were either i) uninfected (Control), ii) SIV-infected, treated with cART and administered either vehicle (VEH) alone (VEH/SIV/cART), or iii) SIV-infected, treated with cART and administered THC:CBD (THC-CBD/SIV/cART). The m^6^A analyses were performed by microarray hybridization profiling following pull down of m^6^A modified miRNAs using an m^6^A specific antibody (Synaptic Systems). BG tissues from all three groups were analyzed for miRNA m^6^A modifications. After quality control and filtering, we used a p-value (unpaired t-test) <= 1.0 and fold-change >= 0.0 to identify a total of 1805 miRNAs that were m^6^A modified in all experimental groups irrespective of statistical significance (**Supplemental Table 1**). We compared each treatment group to the uninfected control group and found that SIV infection induced a significantly broader m^6^A hypomethylation of BG miRNAs (**Table 2**) compared to hypermethylation. Of the total 1805 miRNAs in the VEH/SIV/cART group with m^6^A modifications, 1281 (71%) were mature miRNAs, while 524 (29%) were passenger strand (PS) miRNAs (generated from the opposite (3’) strand) (**Fig. 1A**). Over 95% of the mature and PS miRNAs were m^6^A hypomethylated in VEH/SIV/cART (**Fig. 1B**) and in THC:CBD/SIV/cART (**Fig. 1C**) RMs compared to uninfected controls (**Figs. 1B, 1C**). In contrast, only 6% and 5% of mature and PS miRNAs, respectively, were hypomethylated when comparing VEH/SIV/cART to THC:CBD/SIV/cART (**Fig. 1D**) RMs. Raw intensities of IP of m^6^A-and unmodified-miRNA species, respectively, were normalized with average of log2-scaled spike-in RNA intensities. We found an overall inverse relationship between m^6^A methylation and expression abundance of the miRNA species in VEH/SIV/cART (**Fig. 1E**) and THC:CBD/SIV/cART (**Fig. 1F**) RMs when compared to controls, and that of VEH/SIV/cART when compared to THC:CBD/SIV/cART (**Fig. 1G**) RMs. Our data presented above show that while unsupervised clustering of the differentially expressed miRNAs species isolated from BG fails to separate the control from the THC:CBD treated group, analyses of the differentially m^6^A-modified miRNA clearly identifies changes in the miRNA profile promoted by the THC:CBD treatments, underscoring the need for functional quantification of the RNA levels in addition to the canonical up and down quantifications.

**Figure 1:**
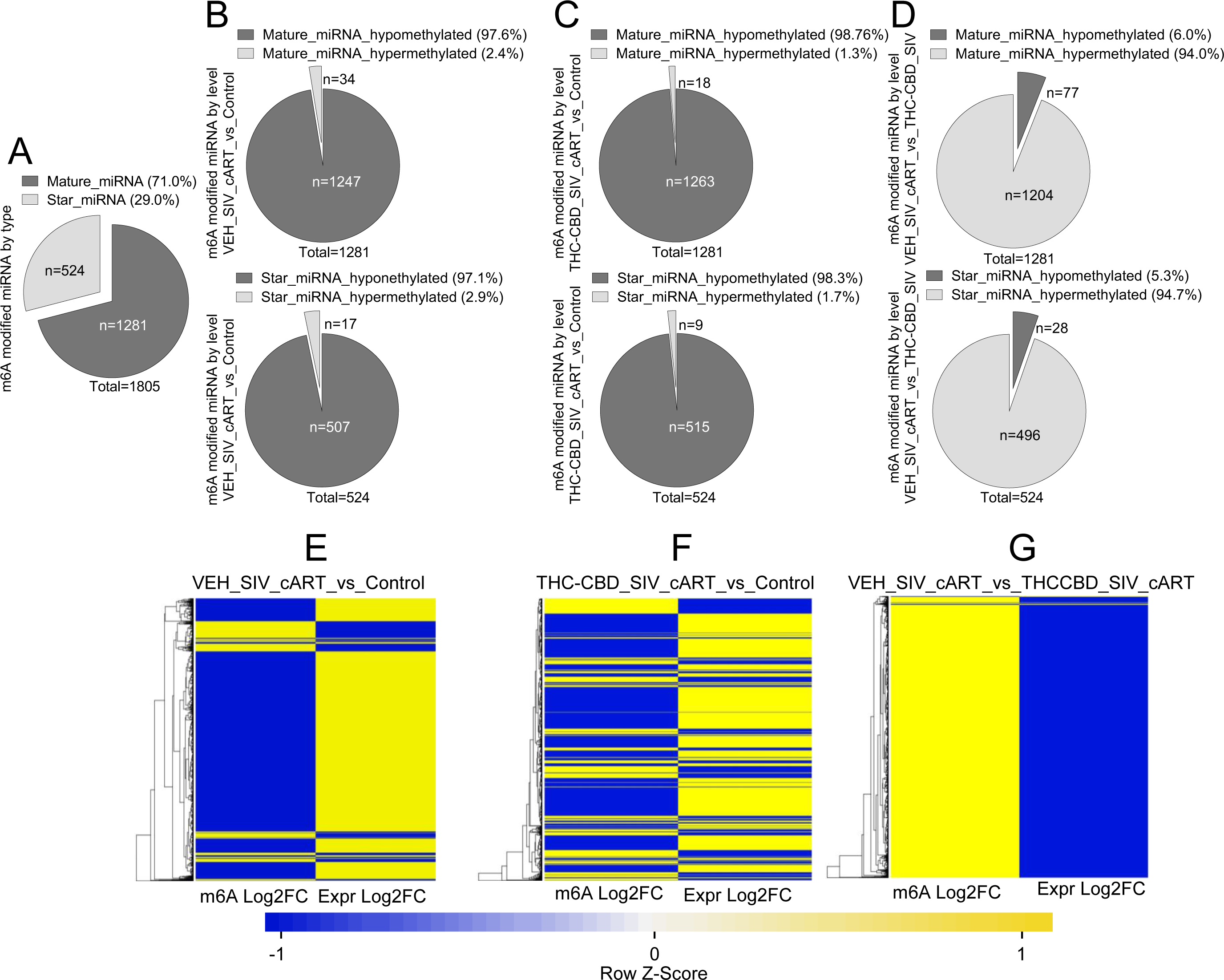
Global m^6^A methylation patterns basal ganglia miRNAs of RM infected with SIV and treated with THC, cART, or both THC and cART over uninfected controls: A) Percent mature and star miRNAs with m^6^A methylation marks. B-D) m^6^A methylation patterns in each treatment group. E-G) Heatmap of m6A methylation profile versus miRNA abundance.

**Table 2:** Total number of identified m^6^A methylated miRNAs per group.

| Table 2: Total number of identified m<sup>6</sup>A methylated miRNAs per group |  |  |  |
| --- | --- | --- | --- |
| Infection and treatment groups | Hypomethylated | Hypermethylated | Total modifications |
| VEH_SIV_cART_vs_Control | 1755 | 50 | **1805** |
| THC-CBD_SIV_cART_vs_Control | 1779 | 26 | **1805** |
| VEH_SIV_cART_vs_THC-CBD_SIV | 105 | 1700 | **1805** |

### SIV infection increased the overall m^6^A methylation levels in BG-derived miRNAs

Further analyses of the m^6^A methylated miRNAs were conducted to identify significantly m^6^A modified miRNAs using p-value < 0.05 (unpaired t-test) and fold change >= 1.5. We found that most of the m^6^A modified miRNAs were present in mature miRNAs with percentages ranging from 60 to 72% (**Figs. 2A-F**). While SIV infection induced significant hypo m^6^A modifications in BG miRNAs in over 1000 miRNA species when compared to control animals, hypermethylation changes were identified in only seven miRNA species (**Table 3**). We next compared methylation levels with miRNA abundance (expression), by calculating the miRNA abundance using mean m^6^A RNA modification quantity per treatment, mean m^6^A RNA modification quantity for control, and the difference of mean percentage of m^6^A RNA modification (%Modified) between the treatment and control groups. The levels of the m^6^A modification versus abundance are displayed as heatmaps (**Figs. 2G-L**). The top 5 miRNAs or fewer (depending on the group) were selected and their m^6^A and expression levels are presented as heatmaps (**Figs. 2M-R**).

**Figure 2:**
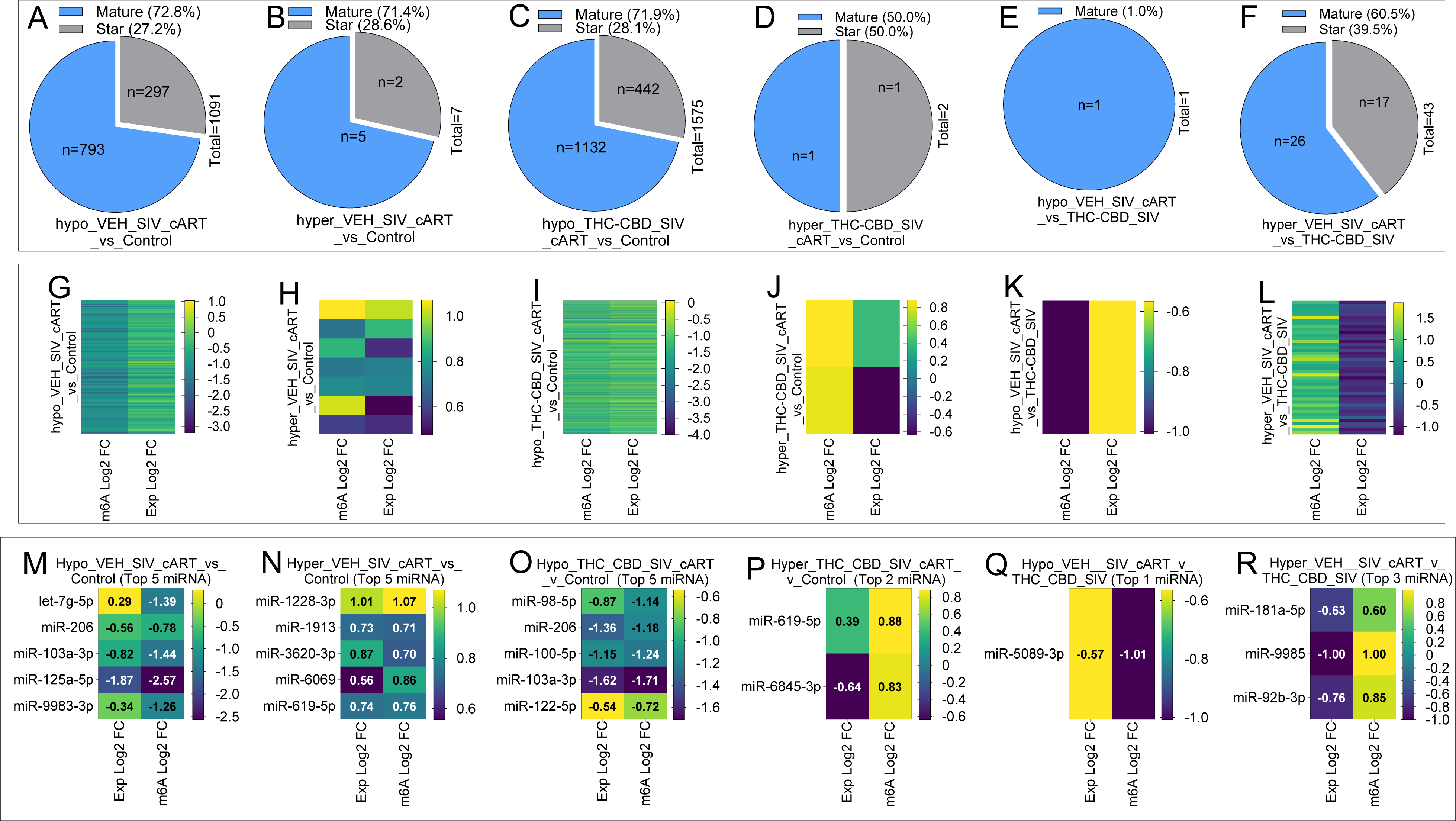
The m6A Content of RM basal ganglia tissues was changed by SIV and THC:CBD: A-F) Pie chart representing % of mature and star miRNAs with m^6^A modifications. G-L) Heatmap illustrating the levels of the m^6^A methylated miRNAs and their abundance. M-R) Heatmap of Top 5 miRNAs from panels G-L.

**Table 3:** Number of signi*ficantly* m^6^A methylated miRNAs per group.

| **Table 3: Number of significantly m<sup>6</sup>A methylated miRNAs per group** |  |
| --- | --- |
| **Infection and treatment groups** | **# of m<sup>6</sup>A modified miRNA** |
| Hypo_VEH/SIV/cART_vs_Control | 1091 |
| Hyper_VEH/SIV_cART_vs_Control | 7 |
| Hypo_THC-CBD/SIV/cART_vs_Control | 1575 |
| Hyper_THC-CBD/SIV/cART_vs_Control | 2 |
| Hypo_VEH/SIV/cART_vs_THC-CBD/SIV | 1 |
| Hyper_VEH/SIV/cART_vs_THC-CBD/SIV | 43 |

### Pathway analysis of SIV-induced m^6^A epitranscriptomic marks in BG miRNAs of SIV-infected RMs identified top neuroinflammation network genes

To analyze the biological implications of m^6^A-enriched miRNAs, we used ingenuity pathway analysis (IPA) to perform enrichment analysis of predicted and validated genes targeted by hypo or hyper m^6^A methylated miRNAs (**Supplemental Table 1**) using criteria determined apriori. We identified neuroinflammation network genes associated with the miRNAs from each group that were hypomethylated (**Supplemental Fig. 1 left, Fig. 3A**) or hypermethylated (**Supplemental Fig. 1 right, Fig. 3B**). The top 35 IPA neuroinflammation network genes associated with m^6^A hypo and hyper methylated miRNAs are shown in **Table 4**. In addition to

**Figure 3:**
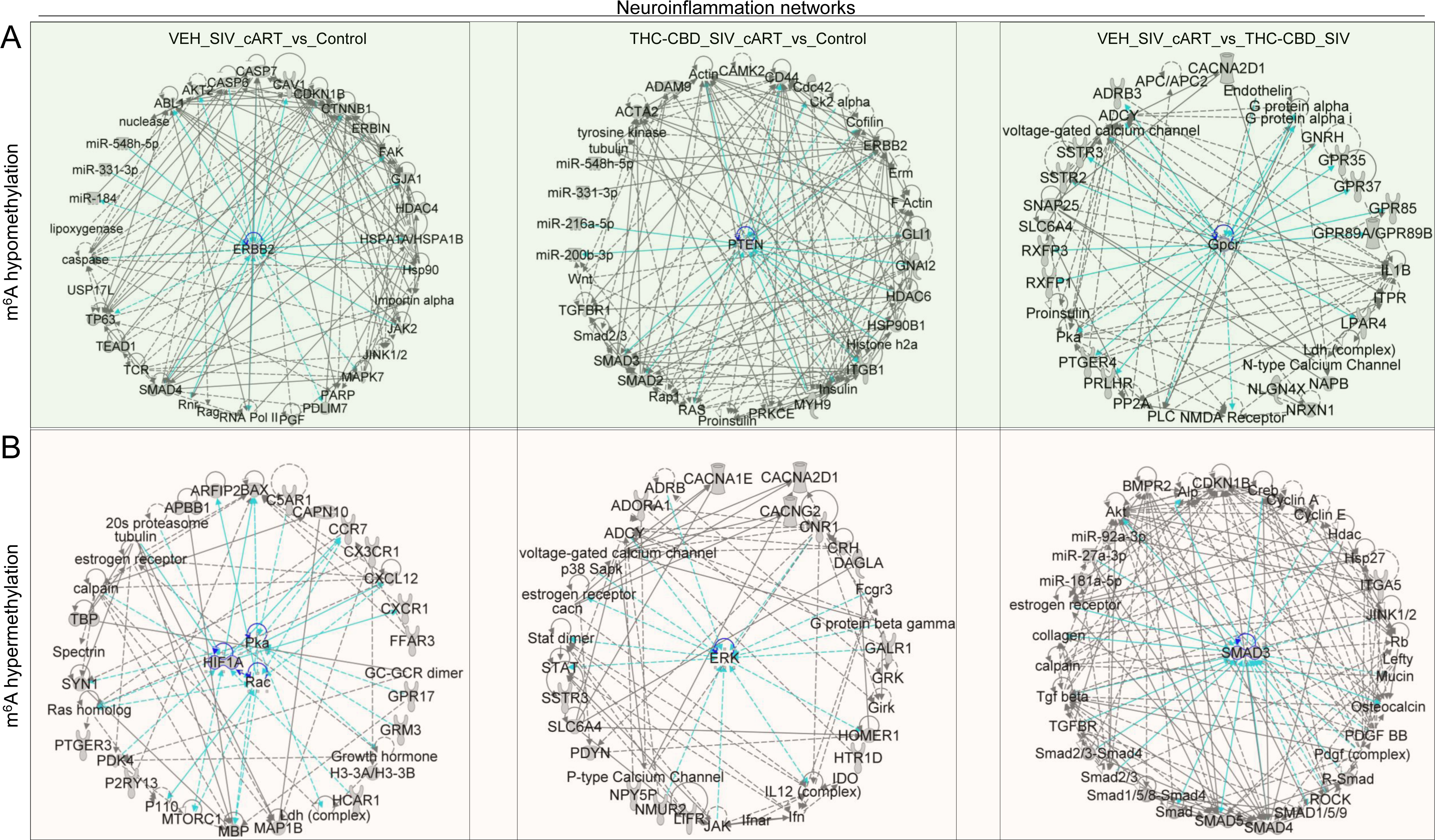
IPA Pathway analysis of m^6^A methylation of basal ganglia miRNAs: A) Neuroinflammation network molecules associated with the with miRNAs with m^6^A hypomethylation (left) or hypermethylation (right). B) Two-way Venny of top 35 network molecules between the hypomethylated and hypermethylated miRNA within each animal groups. C) Three-way Venny of top 30 network molecules comparing m^6^A hypomethylated versus hypermethylated miRNA irrespective of animal groups.

**Table 4:**
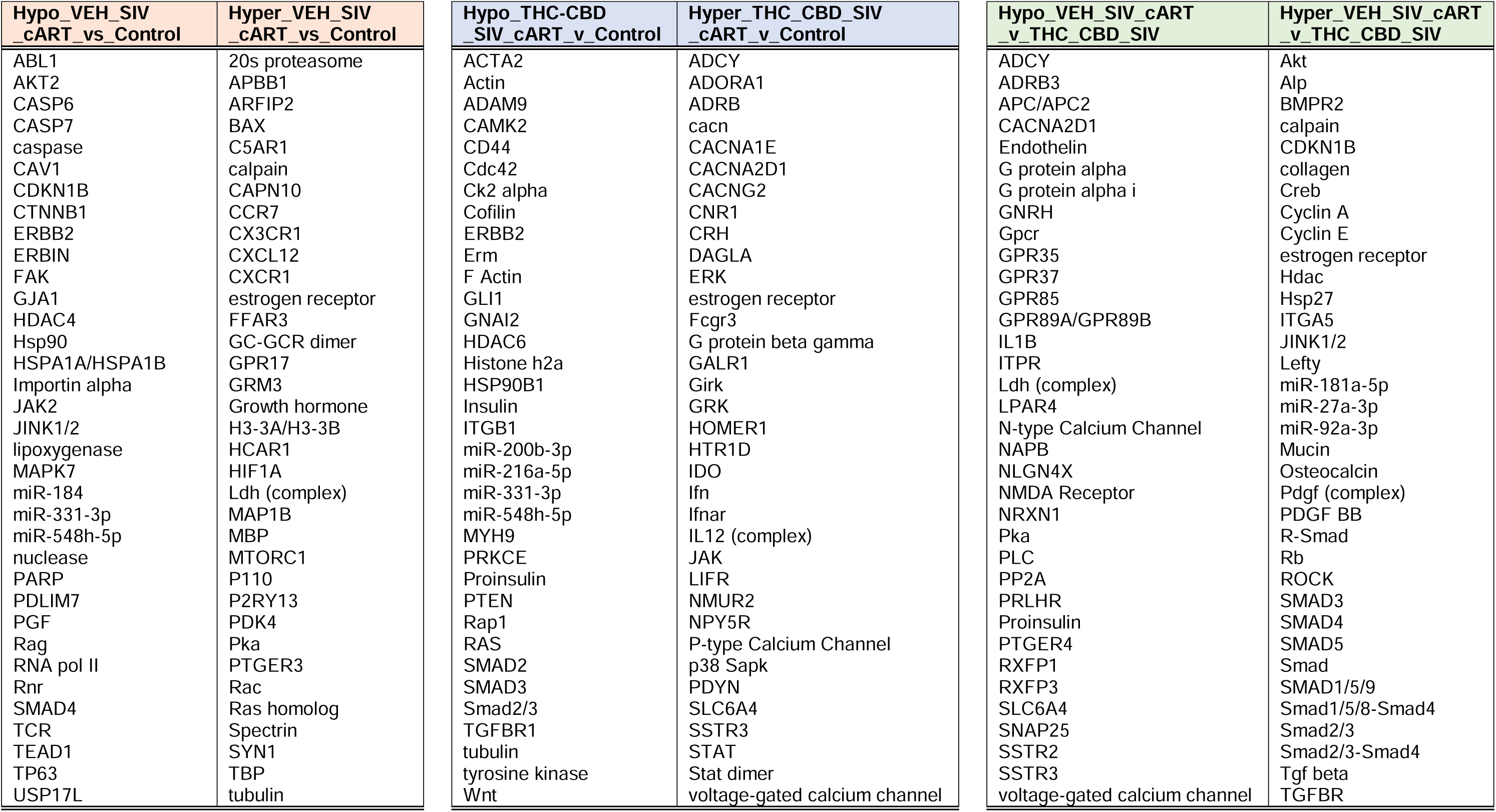
Top IPA neuroinflammation network molecules.

the protein networks altered by the m^6^A methylated miRNAs, our results also identified several miRNA species which were also targeted by the m^6^A-modified miRNAs. These data are consistent with previous reports that demonstrated regulation of primary miR-21 by mature miR-122 [23–25].

### Integration of IPA and GO analyses

Our analyses identified several disease-related functional clusters linked to neuroinflammation network genes associated with m^6^A hypo and hyper methylated miRNAs (**Table 5**). Whereas the top three diseases related to changes in invasion, migration, and proliferation of cells, were linked to a hypomethylated m^6^A profile (rows 1-4), diseases associated with changes in quantity of cells, organization of cytoskeleton, and neurotransmission were linked to the hypermethylated m^6^A profile (rows 1-3). Integration of IPA and the 35 neuroinflammation genes with WEB-based Gene SeT Analysis Toolkit (WebGestalt) GO analysis identified several biological processes, cellular components, and molecular functions of the neuroinflammation network genes for each group that stratified into targets of hypomethylated m^6^A (**Fig. 4A**) and hypermethylated m^6^A (**Fig. 4B**).

**Figure 4:**
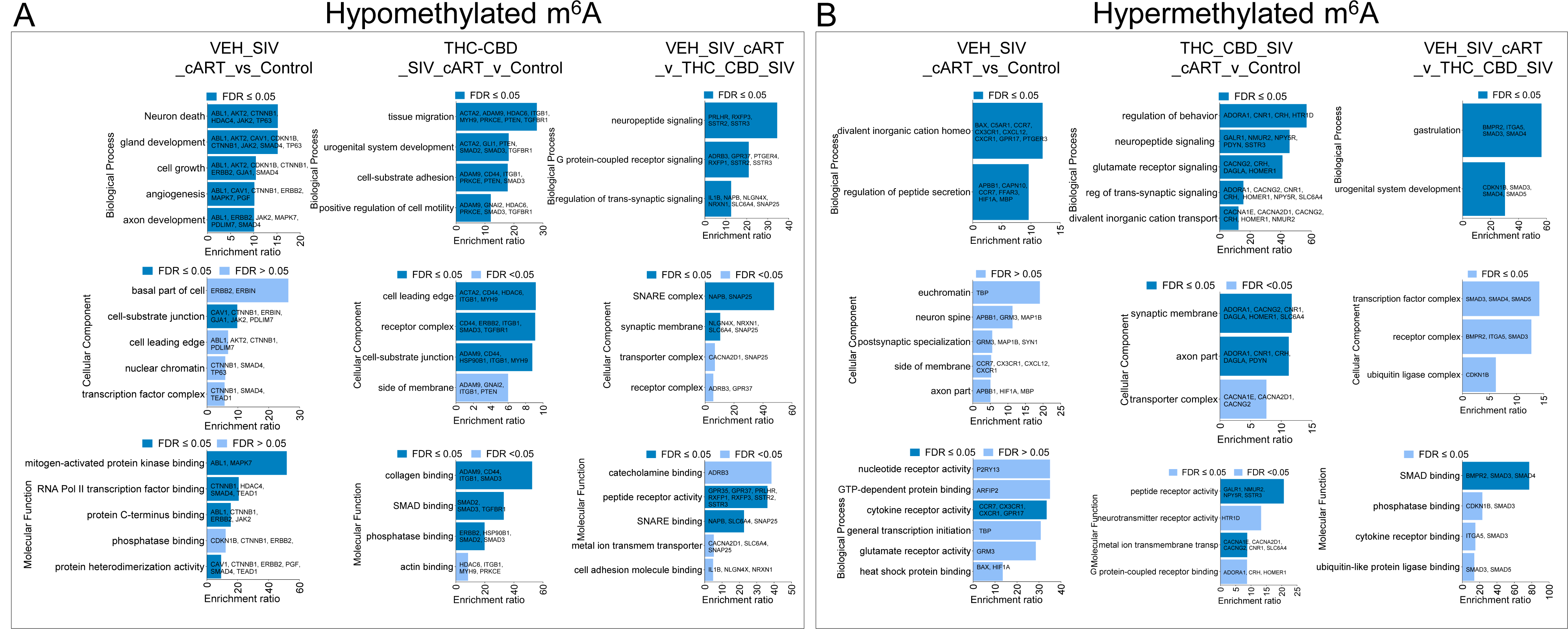
Gene ontology (GO) analysis illustrating biological processes, cellular components, and molecular functions of the neuroinflammation network molecules for each animal with m^6^A: A) hypomethylated and B) hyper methylated miRNA.

**Table 5:** Top Diseases linked to neuroinflammation network molecules.

| Hypo_VEH_SIV_cART_vs_Control |  | Hypo_THC_CBD_SIV_cART_v_Control |  | Hypo_VEH_SIV_cART_v_THC_CBD_SIV |  |
| --- | --- | --- | --- | --- | --- |
| Diseases | p-value | Diseases | p-value | Diseases | p-value |
| Invasion of cells | 1.49E-76 | Invasion of cells | 2.67E-89 | Weight gain | 2.78E-19 |
| Cell movement of tumor cell lines | 7.53E-73 | Cell movement of tumor cell lines | 8.89E-82 | Non-malignant disorder | 2.88E-18 |
| Migration of cells | 1.09E-70 | Cell proliferation of tumor cell lines | 5.38E-80 | Morphology of nervous system | 7.9E-17 |
| Cell proliferation of tumor cell lines | 1.33E-69 | Migration of cells | 5.74E-78 | Bleeding | 8.47E-17 |
| Migration of tumor cell lines | 7.28E-68 | Cell movement | 5.06E-77 | Necrosis | 1.13E-16 |
| Cell movement | 2.81E-67 | Advanced malignant tumor | 5.59E-77 | Hypertrophy | 1.4E-16 |
| Metastatic solid tumor | 3.2E-66 | Advanced malignant solid tumor | 6.3E-77 | Growth of epithelial tissue | 4.45E-16 |
| Advanced malignant solid tumor | 9.23E-66 | Migration of tumor cell lines | 1.18E-76 | Development of vasculature | 4.71E-16 |
| Invasion of tumor cell lines | 5.93E-65 | Apoptosis of tumor cell lines | 1.21E-76 | Cell death of epithelial cells | 5.99E-16 |
| Apoptosis of tumor cell lines | 1.34E-64 | Invasion of tumor cell lines | 1.34E-76 | Benign lesion | 6.2E-16 |
| Metastasis | 4.89E-64 | Metastasis | 1.8E-76 | Seizure disorder | 1.07E-15 |
| Advanced malignant tumor | 5.12E-64 | Metastatic solid tumor | 8.11E-76 | Growth of malignant tumor | 1.17E-15 |
| Cell death of tumor cell lines | 5.15E-63 | Invasive cancer | 2.65E-74 | Necrosis of epithelial tissue | 1.6E-15 |
| Necrosis | 8.22E-63 | Necrosis | 1.23E-73 | Seizures | 2.29E-15 |
| Invasive cancer | 8.5E-63 | Cell death of tumor cell lines | 3.41E-73 | Proliferation of epithelial cells | 6.37E-15 |
| Apoptosis | 5.52E-60 | Apoptosis | 4.18E-69 | Lymphoreticular neoplasm | 9.44E-15 |
| Abnormal morphology of mediastinum | 3.54E-57 | Progressive neurological disorder | 1.22E-66 | Hypertrophy of heart | 1.02E-14 |
| Progressive neurological disorder | 3.74E-56 | Morphology of cardiovascular system | 1.79E-61 | Cognitive impairment | 1.05E-14 |
| Morphology of cardiovascular system | 3.71E-55 | Morphology of heart | 7.78E-61 | Neurological signs | 1.26E-14 |
| Abnormal morphology of heart | 5.19E-55 | Cell survival | 4.1E-60 | Cellular homeostasis | 2.66E-14 |
| Hyper_VEH_SIV_cART_vs_Control |  | Hyper_THC_CBD_SIV_cART_v_Control |  | Hyper_VEH_SIV_cART_v_THC_CBD_SIV |  |
| Diseases | p-value | Diseases | p-value | Diseases | p-value |
| Quantity of cells | 1.32E-66 | Quantity of cells | 6.56E-35 | Migration of cells | 6.84E-14 |
| Organization of cytoskeleton | 5.14E-62 | Neurotransmission | 2.47E-30 | Advanced malignant solid tumor | 2.19E-12 |
| Morphology of nervous system | 2.59E-61 | Morphology of nervous system | 5.56E-30 | Invasion of cells | 2.2E-12 |
| Microtubule dynamics | 3.83E-60 | Transport of molecule | 6.49E-30 | Angiogenesis | 2.88E-12 |
| Proliferation of neural cells | 1.33E-57 | Sensory disorders | 8.73E-29 | Proliferation of epithelial cell lines | 3.49E-12 |
| Organization of cytoplasm | 3.66E-57 | Activation of cells | 1.49E-28 | Vasculogenesis | 8.54E-12 |
| Development of neural cells | 9.77E-57 | Movement Disorders | 2.68E-28 | Proliferation of progenitor cells | 2.76E-11 |
| Neurotransmission | 1.29E-56 | Organization of cytoskeleton | 3.99E-27 | Advanced malignant tumor | 4.8E-11 |
| Movement Disorders | 1.04E-55 | Development of neurons | 6.49E-27 | Proliferation of hematopoietic progenitor cells | 8.05E-11 |
| Development of neurons | 1.31E-53 | Development of neural cells | 7.72E-27 | Advanced extracranial solid tumor | 2.45E-10 |
| Formation of cellular protrusions | 2.5E-53 | Microtubule dynamics | 1.47E-26 | Abnormal morphology of cardio system | 2.45E-10 |
| Proliferation of neuronal cells | 6.67E-53 | Neuronal cell death | 4.02E-26 | Apoptosis of B-lymphocyte derived cell lines | 2.57E-10 |
| Migration of cells | 1.46E-52 | Damage of nervous system | 4.55E-26 | Hepatocellular carcinoma | 2.76E-10 |
| Cell movement | 1.34E-51 | Proliferation of neural cells | 1.09E-25 | Survival of organism | 2.77E-10 |
| Morphology of nervous tissue | 2.21E-51 | Abnormal morphology of nervous sys | 1.29E-25 | Invasion of tumor cell lines | 3.04E-10 |
| Morphogenesis of nervous tissue | 3.38E-50 | Glucose metabolism disorder | 1.96E-25 | Maturation of connective tissue cells | 3.05E-10 |
| Transport of molecule | 5.54E-50 | Morphology of nervous tissue | 3.99E-25 | Invasive cancer | 4.11E-10 |
| Abnormal morphology of nervous sys | 3.14E-49 | Endocrine pancreatic dysfunction | 4.57E-25 | Connective or soft tissue tumor | 4.53E-10 |
| Neuronal cell death | 1.12E-48 | Quantity of neurons | 7.62E-25 | Metastasis | 5.22E-10 |
| Necrosis | 1.79E-48 | Skin sensation disturbance | 7.77E-25 | Metastatic solid tumor | 5.73E-10 |

### Conjoint Analysis of m^6^A methylation and differentially altered miRNA expression

We next determined the functional significance of m^6^A methylation of miRNAs, by first determining the relative abundance of the m^6^A hypo and hyper methylated miRNAs species. After obtaining the abundance of miRNAs with significant m^6^A modifications, we found that majority of significantly m^6^A-modified miRNAs were either significantly up or down regulated (**Table 6**). We also found a positive correlation between m^6^A methylation and miRNA abundance for Hypo_VEH/SIV/cART_vs_Controls (Pearson r=0.72; p<1e-10) and Hypo_THC:CBD/SIV/cART_vs_Controls (Pearson r=0.84; p<1e^-10^) (**Figs. 5A, 5B)**, respectively. However, we did not identify any additional correlations among other treatment groups (**Supplemental Fig. 2**). To examine the functional relationship of m^6^A methylated miRNAs, we used IPA miRNA Target Filter focused on miRNAs related to neurotransmitter networks and nervous system-centered signaling pathways (**Figs. 5C-H**, **Table 7**). Analysis of protein:protein interactions (PPI) for the IPA identified miRNA targets using STRING: functional protein association networks (https://string-db.org/) led to the identification of several gene clusters and nodes associated with the m^6^A methylation status (hypermethylation vs hypomethylation) and miRNA abundance that may shed light into the potential biological functions of miRNA m^6^A methylation (**Figs. 5I-N**). The potential biological functions of the miRNA targets were further revealed by GO biological processes, molecular functions, cellular components, Kegg/Reactome/Wiki pathways, disease gene associations, and tissue expressions (**Table 8**).

**Figure 5:**
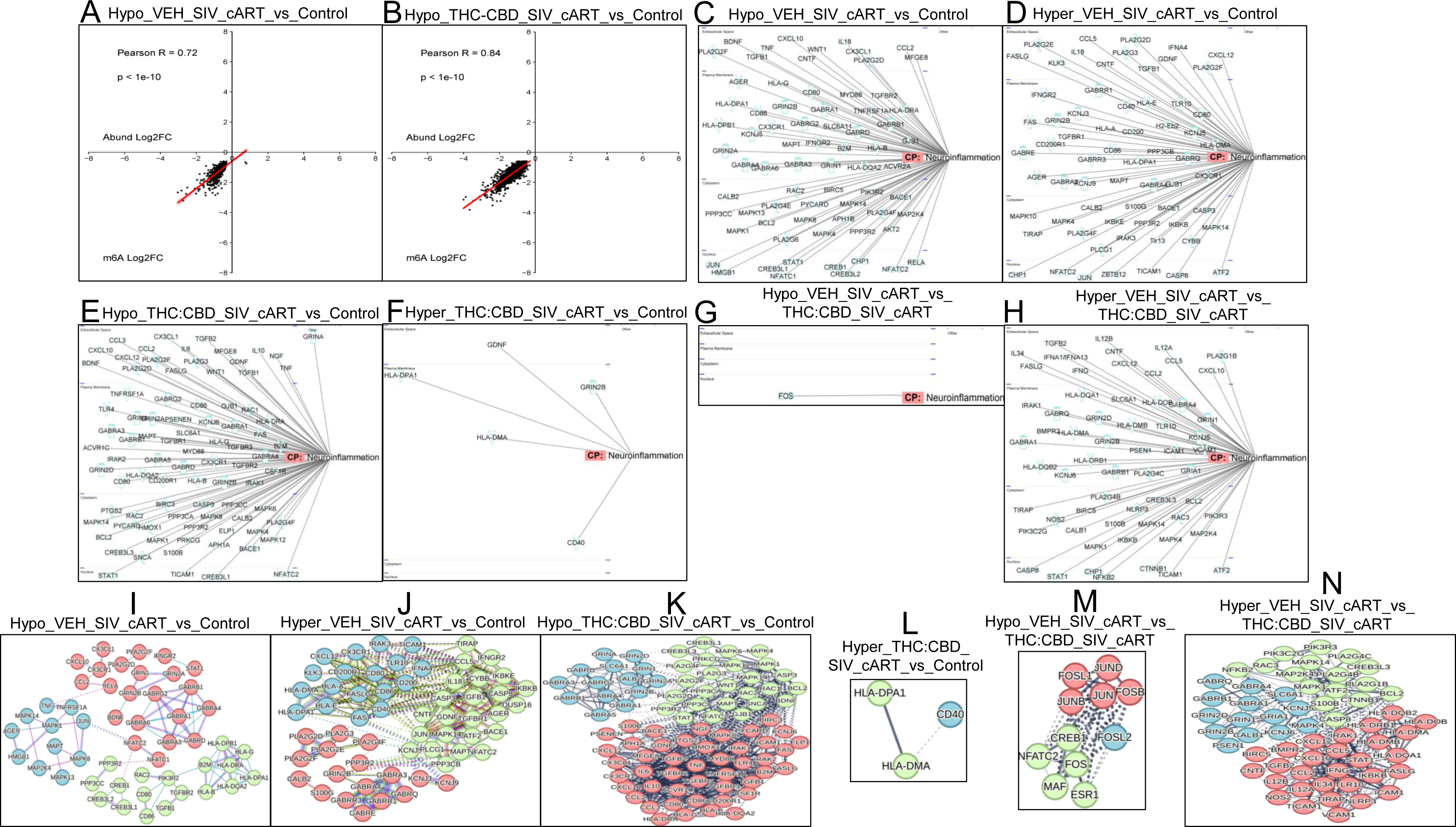
Basal ganglia miRNA with significant m^6^A modifications have altered expression levels. A) Correlation analysis of m^6^A methylation versus miRNA abundance. C-H) Functional relationship of m^6^A methylated miRNA with neurotransmitters/other nervous system and pathogen influenced signaling pathways. I-N) PPI networks for the IPA identified miRNA targets in panels C-H.

**Table 6:**
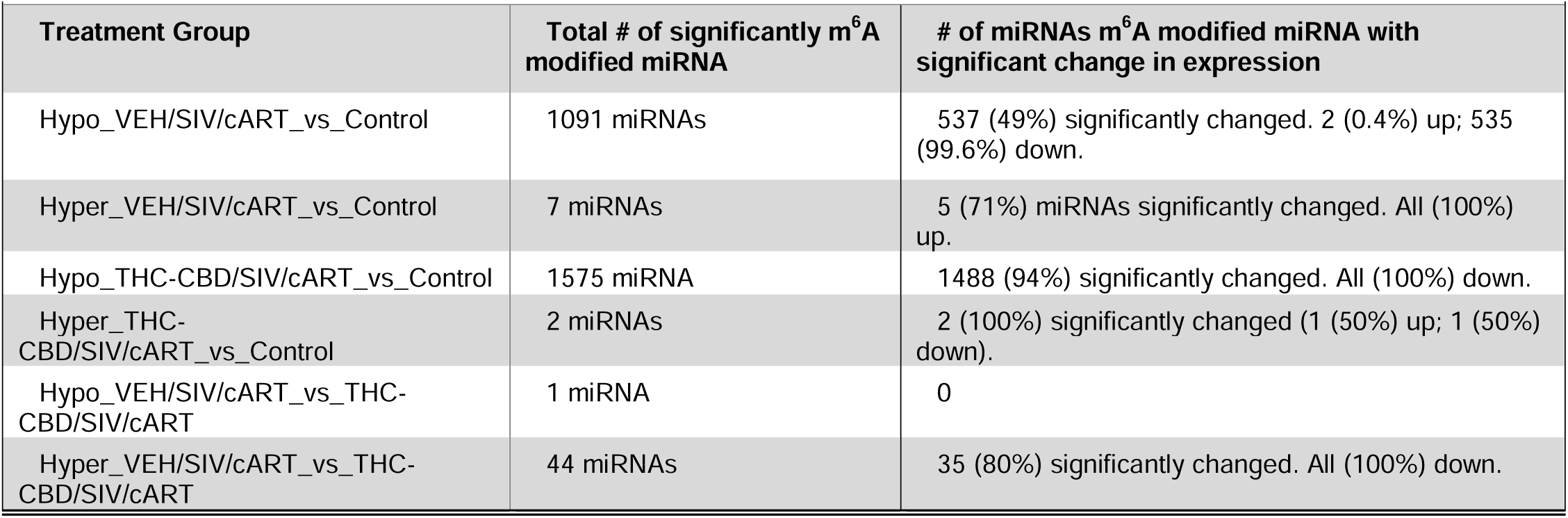
The number of significantly altered miRNAs that were significantly m^6^A modified.

| Treatment Group | Total # of significantly m<sup>6</sup>A modified miRNA | # of miRNAs m<sup>6</sup>A modified miRNA with significant change in expression |
| --- | --- | --- |
| Hypo_VEH/SIV/cART_vs_Control | 1091 miRNAs | 537 (49%) significantly changed. 2 (0.4%) up; 535 (99.6%) down. |
| Hyper_VEH/SIV/cART_vs_Control | 7 miRNAs | 5 (71%) miRNAs significantly changed. All (100%) up. |
| Hypo_THC-CBD/SIV/cART_vs_Control | 1575 miRNA | 1488 (94%) significantly changed. All (100%) down. |
| Hyper_THC-CBD/SIV/cART_vs_Control | 2 miRNAs | 2 (100%) significantly changed (1 (50%) up; 1 (50%) down). |
| Hypo_VEH/SIV/cART_vs_THC-CBD/SIV/cART | 1 miRNA | 0 |
| Hyper_VEH/SIV/cART_vs_THC-CBD/SIV/cART | 44 miRNAs | 35 (80%) significantly changed. All (100%) down. |

**Table 7:**
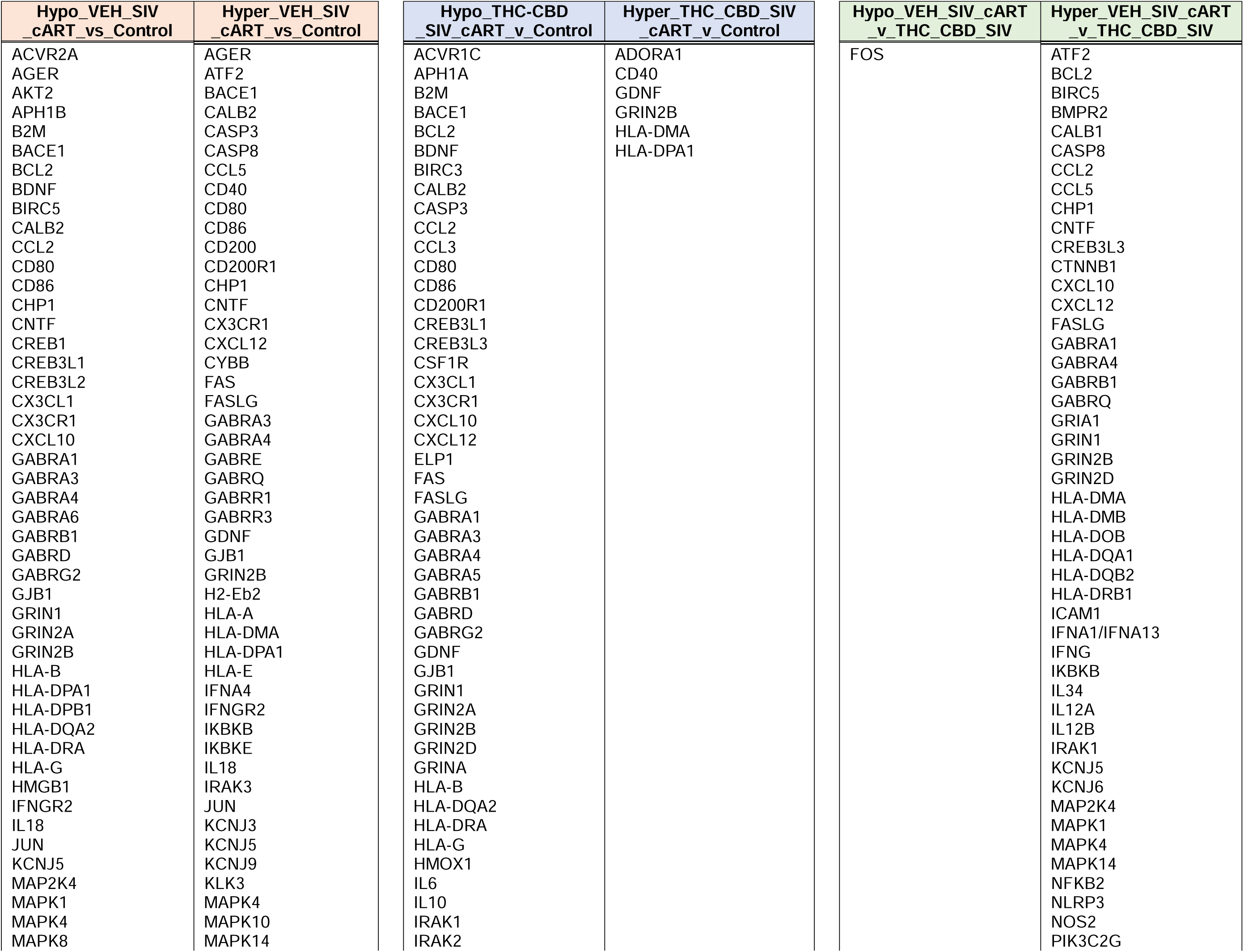
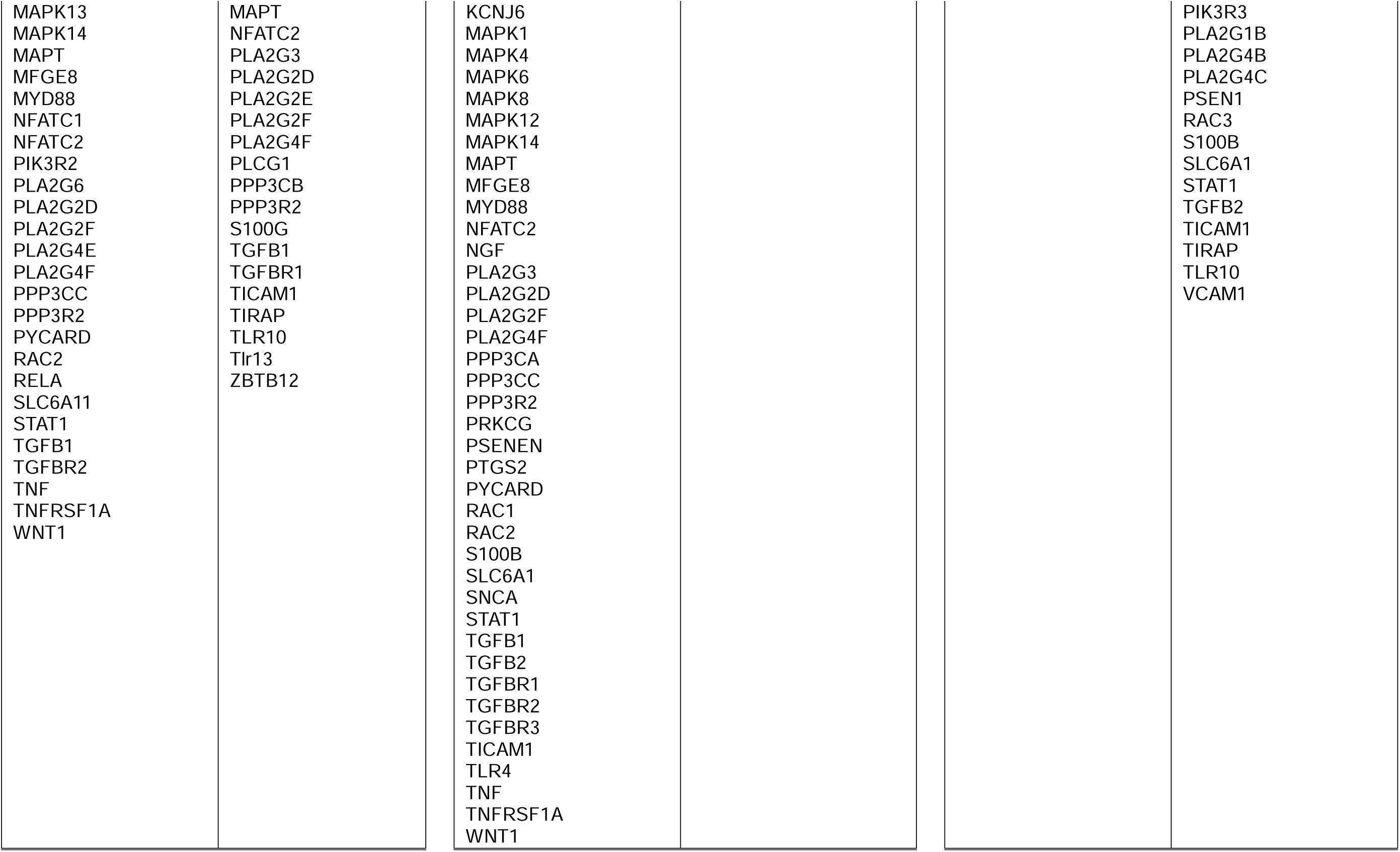
Target genes of miRNAs related to neurotransmitters/other nervous system and pathogen influenced signaling pathways.

**Table 8:**
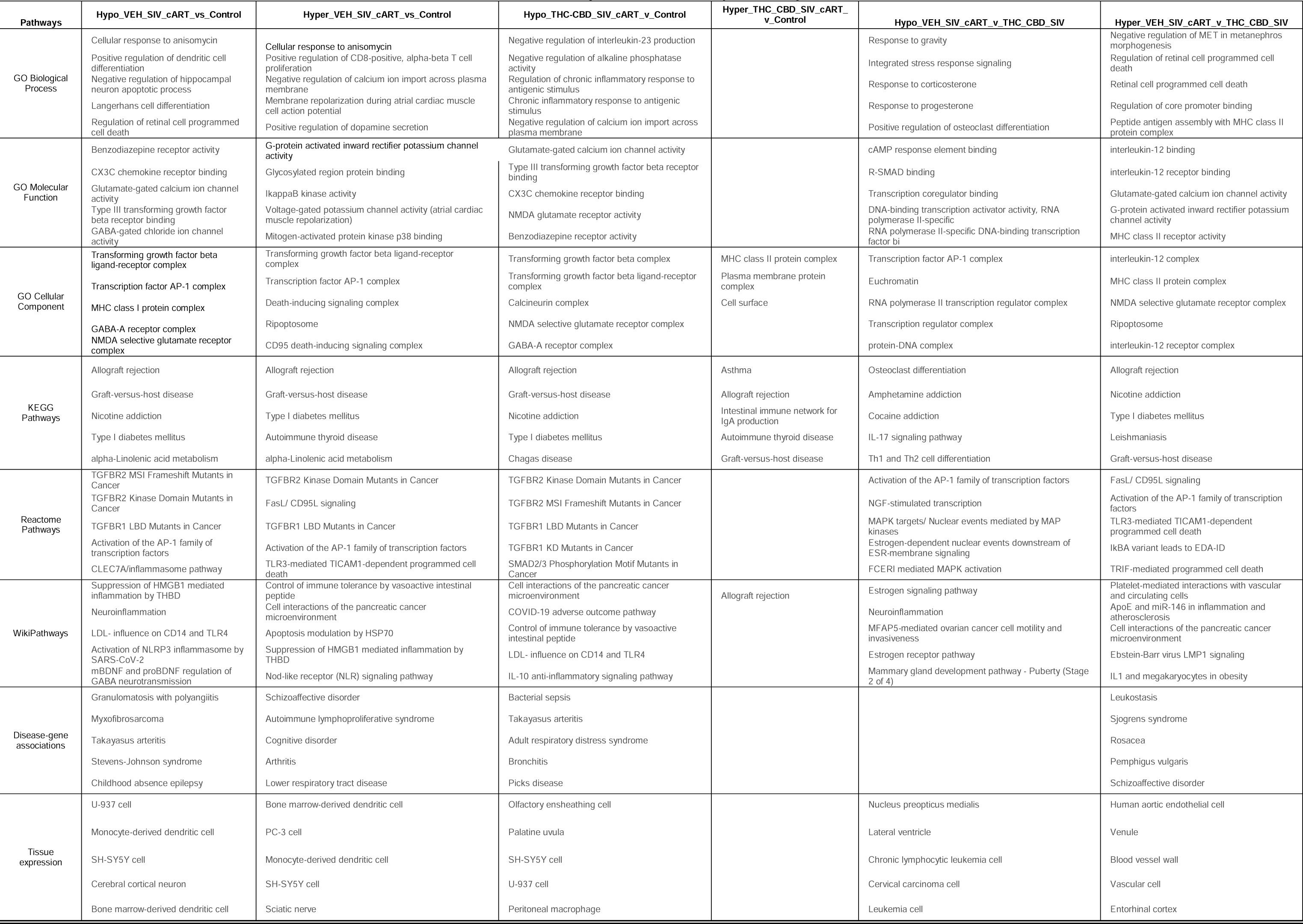
Potential biological functions of the m6A methylated miRNA targets.

### THC:CBD administration significantly reduced the m^6^A levels in the seed sequence of HIV/SIV-altered miRNAs regulating neuroinflammation

Relative to the THC:CBD/SIV/cART group, 44 miRNAs showed significant m^6^A hypermethylation in the VEH/SIV/cART RMs (**Fig. 6A**). More importantly, IPA analyses of the genes targeted by the 44 miRNAs that were hypermethylated in BG of VEH/SIV/cART compared to THC:CBD/SIV/cART RMs (**Table 8**) identified neuroinflammation gene networks previously shown to drive neuroinflammation, neuropathogenesis of Huntington’s, Alzheimer’s, and Parkinson’s disease (**Fig. 6B**). Key biological processes regulated by these m^6^A hypermethylated miRNAs included gamma-aminobutyric acid signaling pathway, synaptic transmission, cognition, immune-response regulating signaling pathway and positive regulation of cytokine production (**Fig. 6C**). GABA receptor activity, glutamate receptor activity, neurotransmitter receptor activity, gated channel activity, cytokine receptor activity were the most significant molecular functions enriched for these mRNA targets. Additionally, the targets of the miRNAs associated with KEGG were linked to GABAergic synapse, leishmaniasis, and nicotine addiction. Reactome pathways are linked to Toll-like receptor 5 cascade, MyD88 cascade, neurotransmitter receptors and postsynaptic signal transmission, signaling by interleukins, and cytokine signaling in the immune system. Diseases associated with autistic disorder, asthma reperfusion injury, atherosclerosis, substance withdrawal syndrome, nerve degeneration, cholestasis, myocardial reperfusion injury, urticaria, hepatic encephalopathy (**Supplemental Figs. 3A-C**. Further analysis of the 44 m^6^A hypermethylated miRNA revealed RRACH or DRACH motif in 14 out of the 44 miRNAs (**Table 9**), indicating that the rest of the m^6^A methylation events took place in non-canonical DRACH motives[26]. IPA network linked the targets of the 14 miRNAs with RRACH or DRACH motif to canonical signaling pathways of NF-kB activation by viruses, neuroinflammation signaling, and transcriptional activity by SMAD heterotrimer, all of which are functionally linked to cognitive impairment[27–29] (**Fig. 6D**).

**Figure 6:**
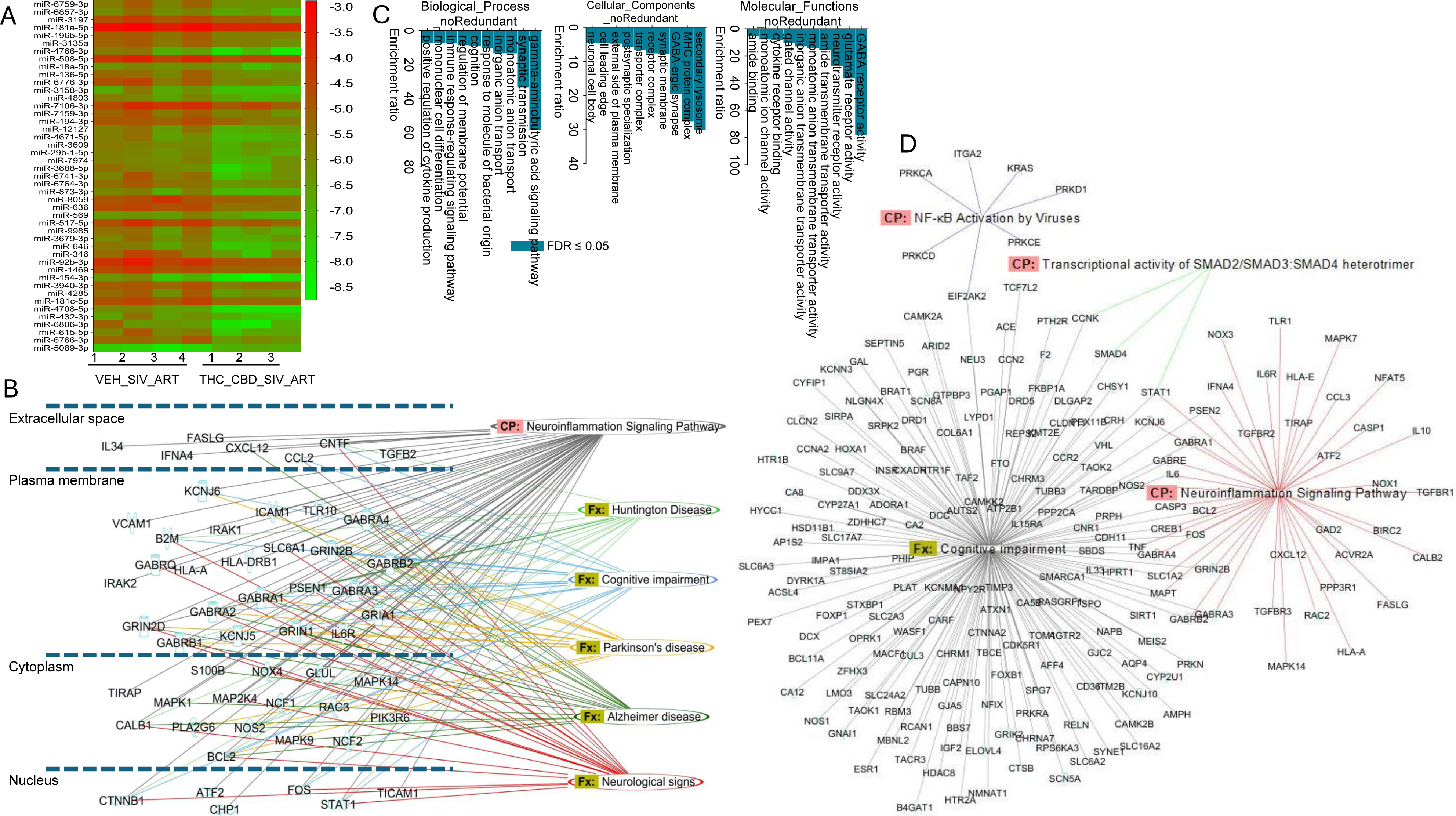
THC:CBD administration significantly reduced the incorporation of m^6^A epitranscriptomic marks on neuroinflammation targeting miRNAs that were increased by HIV/SIV infection: **A)** Heatmap of 44 miRNAs with significant m^6^A hypermethylation in the VEH/SIV/cART RMs relative to THC:CBD/SIV/cART group. **B**) IPA network interactome and associated canonical signaling pathways and biological functions. **C**) Key biological processes, cellular components, and molecular functions regulated by 44 m^6^A hypermethylated miRNAs. **D**) IPA network of canonical signaling pathways and functions associated with 14 miRNAs with RRACH or DRACH motif.

**Table 9:**
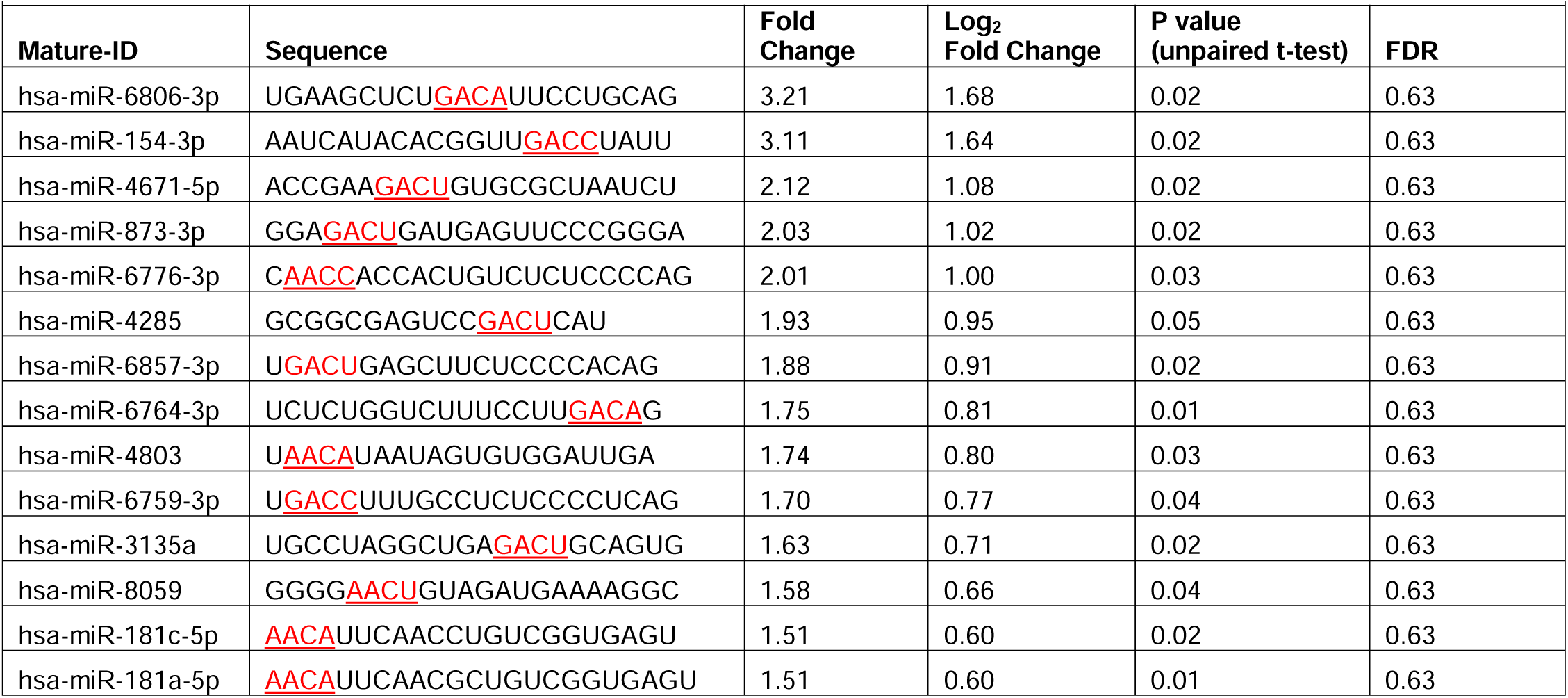
14 miRNAs that were m^6^A hypermethylated bearing RRACH or DRACH motif.

### M6A marks in the miR-194-5p seed region impaired its post-transcriptional silencing potential of the proinflammatory STAT1 protein in vitro

Since miR-194-5p showed significantly more m^6^A hypomethylation in the BG of THC:CBD/SIV/cART (-1.6683 log2 Fold change or -3.178 Fold change) that VE/SIV/cART (-0.8617 log2 Fold change or -1.817 fold change) RMs compared to controls and ∼1.74-fold (p=0.06) more hypomethylation in BG of THC:CBD/SIV/cART relative to VEH/SIV/cART RMs (**Fig. 7B** and **Supplemental Table 1**), we hypothesized that this would allow miR-194-5p to more effectively downregulate its highly conserved (**Fig. 7A**) [30] predicted target gene, STAT1, an interferon-gamma stimulated transcription factor known to activate the expression of interferon-stimulated immune response genes and drive neuroinflammation in PWH and SIV infected RMs [31]. To investigate the impact of m^6^A epitranscriptomic marks on miR-194-5p-STAT1 interactions, we synthesized locked nucleic acid conjugated FAM-labeled m^6^A methylated miR-194-5p (in the seed nucleotide region) mimics along with wild-type miR-194-5p and negative control (NC) mimics and transfected SCC-25 oral squamous cell carcinoma cells. We used SCC-25 cells since they expressed high basal STAT1 protein levels. In contrast, most of the neuronal cells we considered including neuroblastoma cells showed very low to no basal STAT1 protein expression. The miR-194-5p-STAT1 duplex was found to have a minimum free energy of -19.8 kcal/mol (**Fig. 7C**). As evident in **Fig. 7E1** and **E2**, transfection of SCC-2 cells with 30 nM wild type miR-194-5p significantly reduced STAT1 protein expression compared to the wells transfected with negative control (**Fig. 7D1**, **D2** and **G**). Consistent with our hypothesis, adding m^6^A marks to the DRACH motif in the miR-194-5p seed region (**Fig. 7B**) significantly diminished its ability to suppress STAT1 protein expression (**Fig. 7F1**, **F2** and **G**). Accordingly, m6A modified miR-194-5p transfected cells showed significantly higher STAT1 protein expression compared to the cells transfected with wild type miR-194-5p (**Fig. 7F1**, **F2** and **G**) and was similar to the wells transfected with the NC mimic (**Fig. 7D1**, **D2** and **G**).

**Figure 7:**
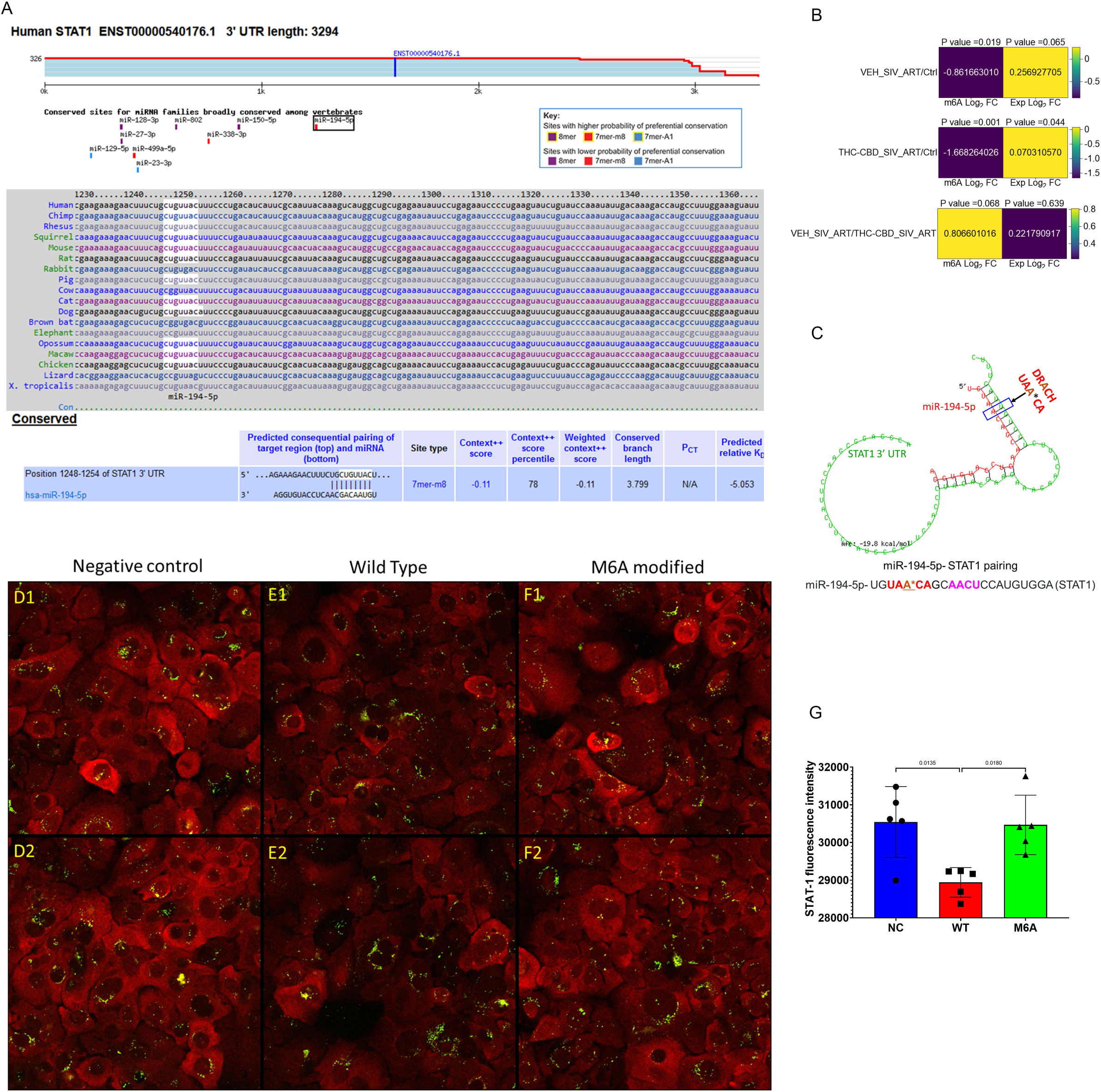
m^6^A marks in the seed region of miR-194-5p impairs its ability to post-transcriptionally silence STAT1 expression. **A)** Conservation of miR-194-5p binding site in the 3’ UTR of STAT1 across twelve different mammalian species identified using TargetScan 8.0. **B)** miR-194-5p fold change in VEH/SIV/cART and THC:CBD/SIV/cART relative to control rhesus macaques. **C)** miR-194-5p-STAT1 pairing as revealed by RNAhybrid has a mean free energy change of -19.8 Kcal/mole. STAT1 protein expression (red) in SCC-25 cells at 72 h post transfection with 30 nM FAM-labeled negative control mimic (**D1** and **D2**), and wild-type (**E1** and **E2**) or m^6^A modified (**F1** and **F2**) (green) miR-194-5p mimics and its quantification using HALO AI software (**G**). Transfections were performed in five triplicate wells for each mimic and experiments were repeated twice. Data were analyzed using One-way ANOVA and post hoc analysis was done using Tukey’s multiple comparison’s test. P<0.05 was considered statistically significant.

### Basal ganglia tissue miRNAs enriched in m^6^A modification are also present in basal ganglia-derived Extracellular Vesicles (EVs)

We integrated BG tissue miRNA abundance with miRNA counts from our previously published BG derived EVs (BG EVs) miRNA sequence dataset [18] to identify tissue miRNAs with m^6^A modifications that were also present in the BG EVs dataset. Of the 527 miRNAs identified in EVs from the BG of SIV infected RMs, 162 (11.1%) were also present in RM whole BG tissues (**Fig. 8A**). Although methylation analyses were not conducted on the EV derived miRNAs, all 162 miRNAs were m^6^A hypomethylated in the BG tissues and none of the m^6^A hypermethylated miRNAs were present in EVs, suggesting either a sorting mechanism based on the methylation status of the miRNAs, or an active de-methylation process performed by m^6^A erasers, which may be present in EVs (**Fig. 8A**). The top 5 EV-associated miRNAs belonged to the let-7 family (let-7a-5p, let-7c-5p, let-7b-5p, miR-26a-5p, let-7f-5p) (**Fig. 8B**) and were different from the top 5 (miR-320b, miR-504-3p, miR-1224-5p, miR-372-3p, miR-193b-5p) miRNAs found in BG tissues (**Fig. 8C**). Next, we used IPA to identify target genes of the 162 BG tissue-EVs miRNAs that were linked to neuroinflammation canonical pathways (**Fig. 8D**). The target genes were used to predict protein-protein interactions to gain insight into potential direct (physical) and indirect (functional) associations. Three gene clusters were identified (**Fig. 8E**). Their predicted GO biological processes detected included processes related to uterine wall breakdown, negative regulation of interleukin-23 production, positive regulation of EMT involved in endocardial cushion formation, positive regulation of tight junction disassembly, positive regulation of additional functions and signaling pathways (**Table 10).** Furthermore, we used WebGestalt to translate the 162 tissue-EV miRNA target gene lists into biological/functional insights using over representation analyses (ORA) with the Benjamini-Hochberg procedure aimed at reducing the false discovery rate and avoiding type I errors. Ten functional categories were enriched, including neuron death, positive regulation of response to external stimulus, leukocyte proliferation, positive regulation of cell motility, regulation of leukocyte, response to molecule of bacterial origin, T cell activation, regulation of cell-cell adhesion, positive regulation of cell activation, extrinsic apoptotic signaling pathway (**Fig. 6F**). The gene list involved in each of the functional categories is presented in **Table 11**.

**Figure 8:**
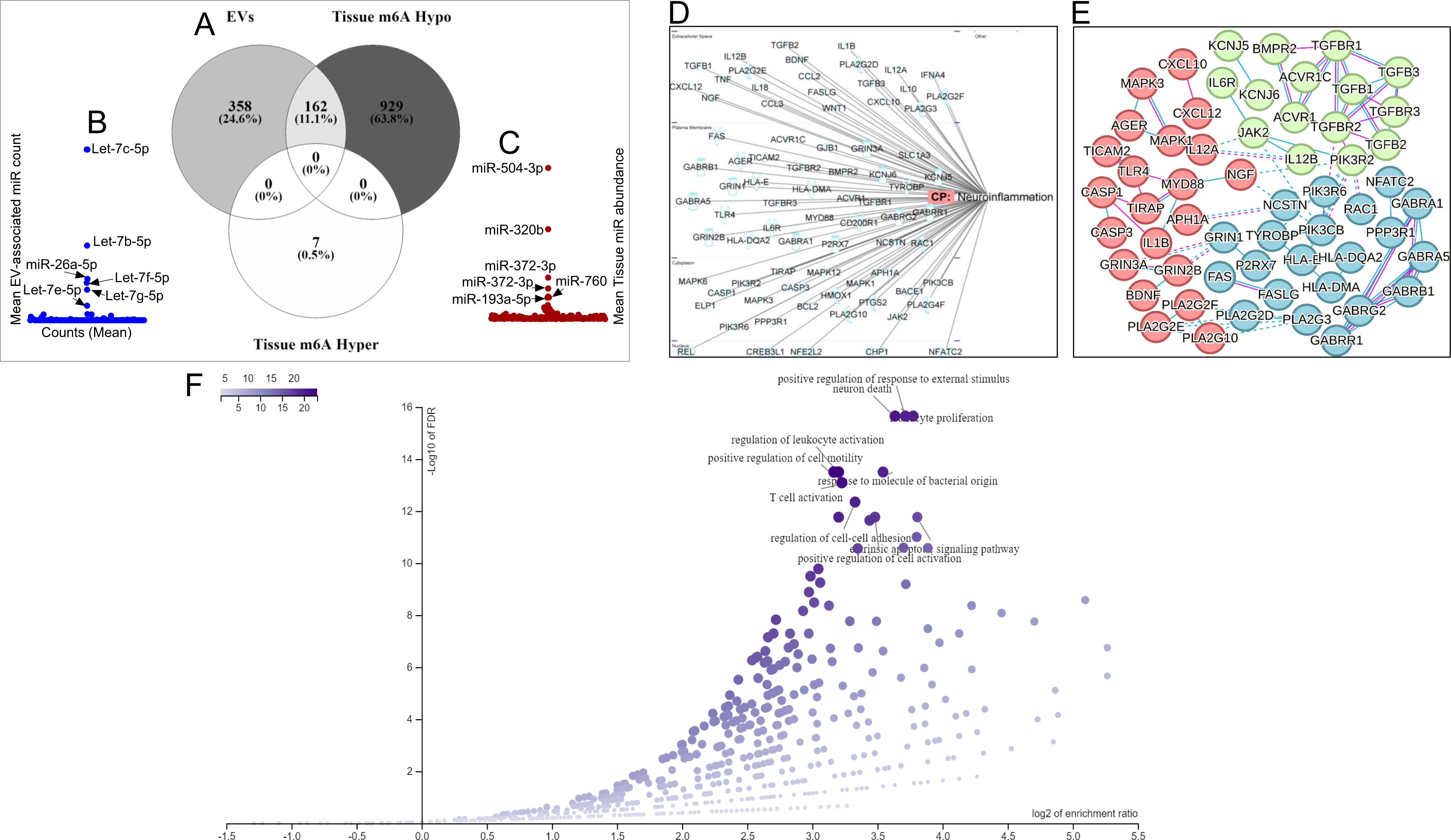
Identification of tissue-EV miRNA with m6A modification: A) 3-way Venn diagram identifying m^6^A modified tissue miRNA present in EVs. B) Top 5 miRNAs in EVs. C) Top 5 miRNAs in tissues. C) Functional relationship of EV-Tissue miRNA with neurotransmitters/other nervous system and pathogen influenced signaling pathways. E) PPI networks for the IPA identified EV-Tissue miRNA targets in panels D. F) Volcano plot of biological/functional insights for EV-Tissue miRNA target gene sets.

**Table 10:**
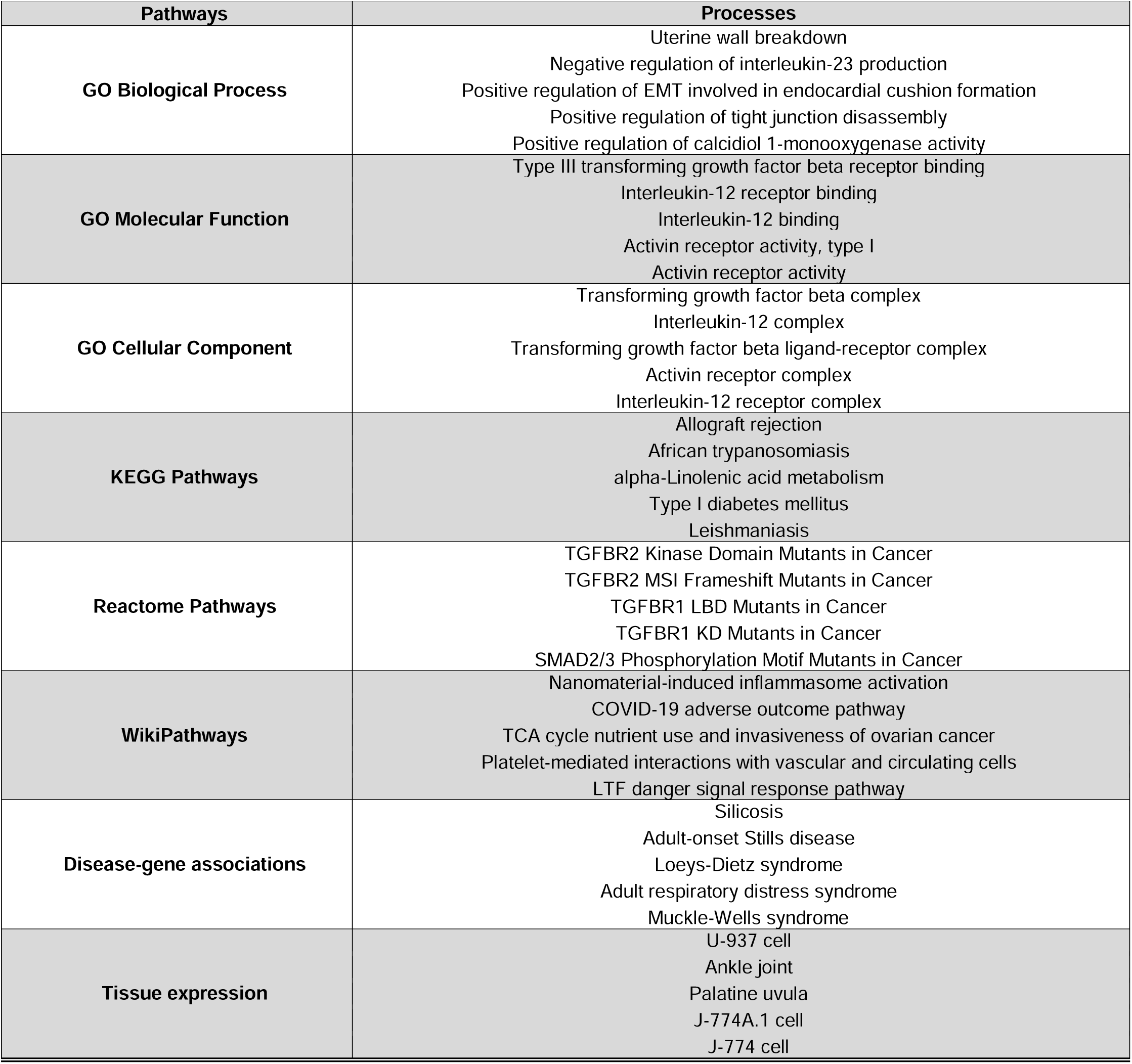
Basal ganglia tissue m^6^A-enriched miRNAs present in Extracellular Vesicles (EVs)

**Table 11:** Gene list involved in functional categories shown in Figure 6F.

| Extrinsic apoptotic signaling pathway | Leukocyte proliferation | Positive regulation of response to external stimulus | Neuron death | Response to molecule of bacterial origin | Positive regulation of cell activation | Regulation of cell-cell adhesion | T cell activation | Regulation of leukocyte activation | Positive regulation of cell motility |
| --- | --- | --- | --- | --- | --- | --- | --- | --- | --- |
| ACVR1 | AGER | AGER | BACE1 | CASP1 | AGER | AGER | AGER | AGER | ACVR1 |
| BCL2 | BCL2 | BMPR2 | BCL2 | CASP3 | BCL2 | CASP3 | BCL2 | BCL2 | BCL2 |
| CASP3 | CASP3 | CCL3 | BDNF | CCL2 | CCL2 | CCL2 | CASP3 | CASP3 | BMPR2 |
| FAS | HLA-E | CXCL10 | CASP3 | CCL3 | HLA-E | CXCL12 | CCL2 | CCL2 | CCL3 |
| FASLG | IFNA4 | CXCL12 | CCL2 | CXCL10 | IL10 | HLA-E | HLA-E | HLA-E | CXCL10 |
| HMOX1 | IL10 | HLA-E | CCL3 | FASLG | IL12A | IL10 | IFNA4 | HMOX1 | CXCL12 |
| IL12A | IL12A | IL12A | CD200R1 | IL10 | IL12B | IL12A | IL10 | IL10 | ELP1 |
| IL1B | IL12B | IL12B | CHP1 | IL12A | IL18 | IL12B | IL12A | IL12A | HMOX1 |
| IL6R | IL18 | IL18 | FASLG | IL12B | IL1B | IL18 | IL12B | IL12B | IL12A |
| JAK2 | IL1B | IL1B | GABRA5 | IL18 | IL6R | IL1B | IL18 | IL18 | IL1B |

## Discussion

Despite the widespread application of multi-omics (mRNA, microRNA and LncRNA profiling) technologies to understanding the pathogenesis of HAND, the field of “epi-transcriptomics” involving post-transcriptional regulation of gene expression via RNA modifications and editing that impacts RNA folding, stability and interaction with its targets remains an underexplored and understudied topic in HIV/SIV infection. Except for a few *in vitro* reports, there exists a dearth of knowledge about m^6^A modifications, the most researched epitranscriptomic modification of eukaryotic RNAs in host–HIV interactions. Moreover, the presence of m^6^A modifications in small RNAs, such as microRNAs (miRNA), and how HIV infection modulates m^6^A modifications in miRNAs during infection is unknown and unexplored. We have previously described the role of miRNAs (both tissue and extracellular vesicle derived) in HIV/SIV induced neuroinflammation and its modulation by long-term low-dose phytocannabinoids. Accordingly, to address this knowledge gap, here, we investigated the profile of BG m^6^A methylome in host miRNAs during HIV/SIV infection and the potential effects of phytocannabinoids (THC and CBD) on the BG miRNA m^6^A methylome. We used a combination of THC and CBD (1:3) because findings from clinical studies have confirmed that THC is well-tolerated when administered in combination with CBD[32–34]. Specifically, CBD diminished the adverse effects of THC, such as tachycardia, intoxication, and sedation, while preserving the beneficial effects of reduced muscle spasticity and neuroinflammation [32–34].

We detected m^6^A modifications of miRNAs from BG tissues of all RMs, irrespective of infection or long-term administration of low-dose phytocannabinoids (THC:CBD). Interestingly, miRNAs from BG tissues of VEH/SIV/cART and THC:CBD/SIV/cART RMs bore significant hypomethylated m^6^A profile compared to uninfected control (VEH) RMs with predictive link to neuroinflammation network genes. The potential functional impact of m^6^A methylation on miRNA function became clearer when we performed a pathway analysis (IPA) of the genes targeted by the 44 miRNAs that were hypermethylated in BG of VEH/SIV/cART compared to THC:CBD/SIV/cART RMs (**Table 2)**. Hypermethylation of miRNA adenine nucleotides in or near the miRNA seed region (nucleotide positions 2 to 7 in the 5’ seed region) can lead to loss of target mRNA 3’ UTR binding and repression[35]. Therefore, in a neuroinflammatory disease like HIV/SIV infection, miRNA m^6^A hypermethylation could lead to enhanced proinflammatory gene expression and disease progression. IPA processing of the target genes identified numerous neuroinflammatory genes such as STAT1, CCL2, CXCL12, IFNA4, TIRPA, NOS2, IL6R, IL34, all known to promote neuroinflammation and drive progression of numerous neurological diseases such as Huntington’s, Alzheimer’s, and Parkinson’s disease, (**Fig 5**). Interestingly, others[31] and we[17] have also shown significantly increased expression of type-I IFN stimulated genes in the brains of SIV-infected RMs. To the best of our knowledge, we provide the first evidence of the presence of m^6^A modifications with a shift towards significant m^6^A hypomethylation in mature miRNAs in the brain during HIV/SIV infection.

The presence of m^6^A modified mature miRNAs was recently reported in HEK293T[36], human lung cancer cells, and fibroblast cells in response to hypoxia[35, 37]. Here, using the SIV-infected RM model, we profiled and identified differential m^6^A methylation in both mature and PS miRNAs in whole BG tissues, although majority of the m^6^A modified miRNAs detected originated from the mature strand. While SIV infection triggered a significant hypomethylated profile in BG tissue miRNAs, miRNA expression abundance was inversely correlated to m^6^A methylation in VEH/SIV/cART and THC:CBD/SIV/cART RMs compared to controls RMs. We also identified a set of BG tissue miRNAs [(n=162) ∼11.1%] with m^6^A epitranscriptomic marks that were also present in BG EVs[18] (**Figure 6**). All 162 BG tissue-EV miRNAs that bore m^6^A marks were hypomethylated and none were hypermethylated.

Next, we used IPA and WebGestalt to predict the potential functional relevance of m^6^A modified miRNAs. Interestingly, miRNAs bearing m^6^A marks were found to have a link to neuroinflammation network genes, neurotransmitters/other nervous system, and pathogen influenced signaling pathways. The top three functional classifications included invasion, migration, and proliferation of cells in generalized disease states linked to m^6^A hypomethylation profile while quantity of cells, organization of cytoskeleton, neurotransmission were the important functional groups/processes found to be linked to the hypermethylated m^6^A profile (**Table 4**). PPI predicted functions and signaling pathways of BG tissue m^6^A-enriched miRNAs that were also present in BG EVs, revealed GO biological processes that included negative regulation of interleukin-23 production, positive regulation of EMT involved in endocardial cushion formation, positive regulation of tight junction disassembly, and positive regulation of calcidiol 1-monooxygenase activity (**Table 8**), all with potential relevance to HIV/SIV induced neurological disease. Recent studies have shown that m^6^A hypermethylation in the seed regions of mature miRNAs resulted in loss of target mRNA recognition, binding, and translational inhibition[35]. Accordingly, based on these predicted functions, it is plausible that the generalized m^6^A hypomethylation may be a host response to enable m^6^A hypomethylated miRNAs to successfully regulate the expression of proinflammatory mediators, such as IL-23, STAT1, NOS1, and the blood brain barrier integrity by promoting posttranscriptional gene silencing.

Although SIV infection decreased m^6^A modifications of BG tissue miRNAs, Lichinchi et al., found that in T cells, HIV infection increased m^6^A marks in both host and viral mRNAs [38]. Altogether, our findings show that BG miRNA m^6^A modifications may play important roles in host–HIV interactions, as previously suggested in other viral infections, although those studies focused entirely on mRNAs [38–40] and not miRNAs.

From a functional perspective, m^6^A miRNA modifications in BG tissues may represent a potential mechanism to regulate host miRNA dependent regulation of proinflammatory gene expression that can directly impact HIV neuropathogenesis. This is evident from recent studies showing m^6^A methylation of the adenine residue/s in miR-21-5p[35], where m^6^A modification of adenine residues near but not in the seed region resulted in loss of target gene (MAPK10 and PTEN) repression in A549 cells. Therefore, the significantly reduced m^6^A methylation in 44 neuroinflammatory gene targeting miRNAs in the THC:CBD/SIV/cART compared to the VEH/SIV/cART group suggests that cannabinoids may preserve the post-transcriptional gene regulatory functions of anti-inflammatory miRNAs by reducing their m^6^A epitranscriptomic marks. Our recently published studies found long-term low-dose THC administration to significantly reduce type I-interferon stimulated gene expression[17]. Since type-I IFNs activate STAT1[41], we performed bioinformatics analysis using TargetScan 8.0 and identified RM STAT1 to be a predicted target of miR-194-5p with a perfect match between the seed sequence of miR-194-5p and the 3’ UTR region of STAT1 (**Fig. 7A**). Interestingly, miR-194-5p showed markedly more hypomethylation in THC:CBD/SIV/cART (-1.6683 log2 Fold change) than VEH/SIV/cART (-0.8617 log2 Fold change) compared to control RMs. Increased STAT1 activated by interferons (IFN) is known to drive persistent neuroinflammatory signaling in a variety of neurological diseases such as Alzheimer’s, Parkinson’s, Huntington’s disease and traumatic brain injury. In addition, significant upregulation of IFN-stimulated STAT1 protein expression in the brain has been confirmed by others in the SIV-infected RM model[31]. We further identified a DRACH motif (where, D= A, G or U; U, R= A or G, and H=A, C or U) in the seed region (nucleotides 2-7 from the 5’ end) of miR-194-5p (5’-UG**UAA*CA**GC**AACU**CCAUGUGGA-3’).

Under in vitro conditions, m^6^A methylation of adenine residues in mature miRNAs resulted in reduced suppression of target gene expression[35], suggesting that m^6^A modification of miRNA seed nucleotides can interfere with mRNA target recognition and binding. Consistent with the findings of Wang et al. [35], in vitro overexpression of miR-194-5p mimics containing m^6^A methylated adenosine nucleotides in the seed region failed to downregulate STAT1 protein expression, while wildtype miR-194-5p significantly reduced STAT1 protein expression at 72 h post transfection. Overall, these findings suggest that the presence of m^6^A marks in the miR-194-5p seed region could potentially destabilize A:U pairing, thereby weakening its interactions with its binding region in the 3’ UTR of STAT1 mRNA. On the contrary, reduced m^6^A methylation of miR-194-5p can preserve its target binding capacity and represents a potential posttranscriptional gene silencing mechanism by which THC or THC:CBD may reduce STAT1 expression and accordingly reduce proinflammatory signaling in the brain and periphery. It is possible that THC:CBD may mediate these changes through altering the expression levels of m^6^A writers and erasers that catalyze the addition and removal of m^6^A epitranscriptomic marks.

While we m^6^A marks in the miR-194-5p seed region interfered with its ability to downregulate STAT1 protein expression, it remains to be determined how m^6^A marks in the 3’ and regions adjacent to seed nucleotide impact miRNA function. Therefore, future studies are needed to address these knowledge gaps including whether extracellular miRNAs bear similar m^6^A marks and the implications of such epitranscriptomic marks in HIV neuropathogenesis.

### Declarations

**Consent for publication:** All authors read and approved the publication of this manuscript.

**Competing interests:** The authors report no biomedical financial interests or potential conflicts of interest.

**Availability of data and materials:** The dataset is included within the article and its additional files.

**Funding:** This work was supported by National Institutes of Health funding (Grant No. R01DA042348 [to CMO]; Grant Nos. R01DA042524 and R01 DA052845 [MM], Grant Nos. R01DA050169 and R21/R33DA053643 [to CMO & MM], R01CA218500, UH3TR002881, R01TR218500 [to IG], P30AI161943, and P51OD111033.

**Author contributions:** Study conceptualization and design: IG, CST and MM. CMO conducted data analyses. CMO, MM, and IG wrote the original draft, reviewed and edited the manuscript. IG, LSP and MM performed the in vitro m^6^A miR-194-5p overexpression and immunofluorescence studies and HALO image analysis and quantification. LSP assisted with the viral load quantification, manuscript proof reading and preparation of animal information.

**Supplemental Figure 1: IPA Pathway analysis of m^6^A methylation of basal ganglia miRNAs:** ) Neuroinflammation network molecules associated with the with miRNAs with m6A hypomethylation (left) or hypermethylation (right).

**Supplemental Figure 2: Correlation analysis of m^6^A methylation versus miRNA abundance:**

**Supplemental Figure 3: Web Gestalt analysis for A) KEGG, B) Reactome, C) Diseases.**

## References

1. Eden A, Fuchs D, Hagberg L, Nilsson S, Spudich S, Svennerholm B, Price RW, Gisslen M: HIV-1 viral escape in cerebrospinal fluid of subjects on suppressive antiretroviral treatment. The Journal of infectious diseases 2010, 202:1819–1825.

2. Lamers SL, Gray RR, Salemi M, Huysentruyt LC, McGrath MS: HIV-1 phylogenetic analysis shows HIV-1 transits through the meninges to brain and peripheral tissues. Infection, genetics and evolution : journal of molecular epidemiology and evolutionary genetics in infectious diseases 2011, 11:31–37.

3. Spudich S, Lollo N, Liegler T, Deeks SG, Price RW: Treatment benefit on cerebrospinal fluid HIV-1 levels in the setting of systemic virological suppression and failure. The Journal of infectious diseases 2006, 194:1686–1696.

4. Grant I, Atkinson JH, Hesselink JR, Kennedy CJ, Richman DD, Spector SA, McCutchan JA: Evidence for early central nervous system involvement in the acquired immunodeficiency syndrome (AIDS) and other human immunodeficiency virus (HIV) infections. Studies with neuropsychologic testing and magnetic resonance imaging. Annals of internal medicine 1987, 107:828–836.

5. Sacktor N, Skolasky RL, Seaberg E, Munro C, Becker JT, Martin E, Ragin A, Levine A, Millerp E: Prevalence of HIV-associated neurocognitive disorders in the Multicenter AIDS Cohort Study. Neurology 2016, 86:334–340.

6. Fuster-RuizdeApodaca MJ, Castro-Granell V, Garin N, Laguía A, Jaén Á, Iniesta C, Cenoz S, Galindo MJ: Prevalence and patterns of illicit drug use in people living with HIV in Spain: A cross-sectional study. PLoS One 2019, 14:e0211252.

7. Castro FOF, Silva JM, Dorneles GP, Barros JBS, Ribeiro CB, Noronha I, Barbosa GR, Souza LCS, Guilarde AO, Pereira A, et al: Distinct inflammatory profiles in HIV-infected individuals under antiretroviral therapy using cannabis, cocaine or cannabis plus cocaine. Aids 2019, 33:1831–1842.

8. Galaj E, Bi GH, Yang HJ, Xi ZX: Cannabidiol attenuates the rewarding effects of cocaine in rats by CB2, 5-HT(1A) and TRPV1 receptor mechanisms. Neuropharmacology 2020, 167:107740.

9. Mimiaga MJ, Reisner SL, Grasso C, Crane HM, Safren SA, Kitahata MM, Schumacher JE, Mathews WC, Mayer KH: Substance use among HIV-infected patients engaged in primary care in the United States: findings from the Centers for AIDS Research Network of Integrated Clinical Systems cohort. Am J Public Health 2013, 103:1457–1467.

10. Mohammadi A, Darabi M, Nasry M, Saabet-Jahromi MJ, Malek-Pour-Afshar R, Sheibani H: Effect of opium addiction on lipid profile and atherosclerosis formation in hypercholesterolemic rabbits. Exp Toxicol Pathol 2009, 61:145–149.

11. Roohafza H, Talaei M, Sadeghi M, Haghani P, Shokouh P, Sarrafzadegan N: Opium decreases the age at myocardial infarction and sudden cardiac death: a long- and short-term outcome evaluation. Arch Iran Med 2013, 16:154–160.

12. Nabati S, Asadikaram G, Arababadi MK, Shahabinejad G, Rezaeian M, Mahmoodi M, Kennedy D: The plasma levels of the cytokines in opium-addicts and the effects of opium on the cytokines secretion by their lymphocytes. Immunol Lett 2013, 152:42–46.

13. Saha B, Momen-Heravi F, Kodys K, Szabo G: MicroRNA Cargo of Extracellular Vesicles from Alcohol-exposed Monocytes Signals Naive Monocytes to Differentiate into M2 Macrophages. J Biol Chem 2016, 291:149–159.

14. Momen-Heravi F, Saha B, Kodys K, Catalano D, Satishchandran A, Szabo G: Increased number of circulating exosomes and their microRNA cargos are potential novel biomarkers in alcoholic hepatitis. J Transl Med 2015, 13:261.

15. Momen-Heravi F, Bala S, Kodys K, Szabo G: Exosomes derived from alcohol-treated hepatocytes horizontally transfer liver specific miRNA-122 and sensitize monocytes to LPS. Sci Rep 2015, 5:9991.

16. McDew-White M, Lee E, Alvarez X, Sestak K, Ling BJ, Byrareddy SN, Okeoma CM, Mohan M: Cannabinoid control of gingival immune activation in chronically SIV-infected rhesus macaques involves modulation of the indoleamine-2,3-dioxygenase-1 pathway and salivary microbiome. EBioMedicine 2022, 75:103769.

17. McDew-White M, Lee E, Premadasa LS, Alvarez X, Okeoma CM, Mohan M: Cannabinoids modulate the microbiota-gut-brain axis in HIV/SIV infection by reducing neuroinflammation and dysbiosis while concurrently elevating endocannabinoid and indole-3-propionate levels. J Neuroinflammation 2023, 20:62.

18. Kaddour H, McDew-White M, Madeira MM, Tranquille MA, Tsirka SE, Mohan M, Okeoma CM: Chronic delta-9-tetrahydrocannabinol (THC) treatment counteracts SIV-induced modulation of proinflammatory microRNA cargo in basal ganglia-derived extracellular vesicles. J Neuroinflammation 2022, 19:225.

19. Jonkhout N, Tran J, Smith MA, Schonrock N, Mattick JS, Novoa EM: The RNA modification landscape in human disease. Rna 2017, 23:1754–1769.

20. Fan Y, Lv X, Chen Z, Peng Y, Zhang M: m6A methylation: Critical roles in aging and neurological diseases. Front Mol Neurosci 2023, 16:1102147.

21. Zhang F, Ran Y, Tahir M, Li Z, Wang J, Chen X: Regulation of N6-methyladenosine (m6A) RNA methylation in microglia-mediated inflammation and ischemic stroke. Front Cell Neurosci 2022, 16:955222.

22. Chatterjee B, Shen CJ, Majumder P: RNA Modifications and RNA Metabolism in Neurological Disease Pathogenesis. Int J Mol Sci 2021, 22.

23. Hill M, Tran N: Global miRNA to miRNA Interactions: Impacts for miR-21. Trends Cell Biol 2021, 31:3–5.

24. Ooi JYY, Bernardo BC, Singla S, Patterson NL, Lin RCY, McMullen JR: Identification of miR-34 regulatory networks in settings of disease and antimiR-therapy: Implications for treating cardiac pathology and other diseases. RNA Biol 2017, 14:500–513.

25. Wang D, Sun X, Wei Y, Liang H, Yuan M, Jin F, Chen X, Liu Y, Zhang CY, Li L, Zen K: Nuclear miR-122 directly regulates the biogenesis of cell survival oncomiR miR-21 at the posttranscriptional level. Nucleic Acids Res 2018, 46:2012–2029.

26. Price AM, Hayer KE, McIntyre ABR, Gokhale NS, Abebe JS, Della Fera AN, Mason CE, Horner SM, Wilson AC, Depledge DP, Weitzman MD: Direct RNA sequencing reveals m(6)A modifications on adenovirus RNA are necessary for efficient splicing. Nat Commun 2020, 11:6016.

27. Kashima R, Hata A: The role of TGF-beta superfamily signaling in neurological disorders. Acta Biochim Biophys Sin (Shanghai) 2018, 50:106–120.

28. Su C, Miao J, Guo J: The relationship between TGF-beta1 and cognitive function in the brain. Brain Res Bull 2023, 205:110820.

29. Deng Z, Fan T, Xiao C, Tian H, Zheng Y, Li C, He J: TGF-beta signaling in health, disease, and therapeutics. Signal Transduct Target Ther 2024, 9:61.

30. Agarwal V, Bell GW, Nam JW, Bartel DP: Predicting effective microRNA target sites in mammalian mRNAs. Elife 2015, 4.

31. Roberts ES, Burudi EM, Flynn C, Madden LJ, Roinick KL, Watry DD, Zandonatti MA, Taffe MA, Fox HS: Acute SIV infection of the brain leads to upregulation of IL6 and interferon-regulated genes: expression patterns throughout disease progression and impact on neuroAIDS. J Neuroimmunol 2004, 157:81–92.

32. Cooper DB, Bunner AE, Kennedy JE, Balldin V, Tate DF, Eapen BC, Jaramillo CA: Treatment of persistent post-concussive symptoms after mild traumatic brain injury: a systematic review of cognitive rehabilitation and behavioral health interventions in military service members and veterans. Brain Imaging Behav 2015, 9:403–420.

33. Russo E, Guy GW: A tale of two cannabinoids: the therapeutic rationale for combining tetrahydrocannabinol and cannabidiol. Med Hypotheses 2006, 66:234–246.

34. Vermersch P: Sativex(®) (tetrahydrocannabinol + cannabidiol), an endocannabinoid system modulator: basic features and main clinical data. Expert Rev Neurother 2011, 11:15–19.

35. Wang H, Song X, Song C, Wang X, Cao H: m(6)A-seq analysis of microRNAs reveals that the N6-methyladenosine modification of miR-21-5p affects its target expression. Arch Biochem Biophys 2021, 711:109023.

36. Berulava T, Rahmann S, Rademacher K, Klein-Hitpass L, Horsthemke B: N6-adenosine methylation in MiRNAs. PLoS One 2015, 10:e0118438.

37. van den Homberg DAL, van der Kwast R, Quax PHA, Nossent AY: N-6-Methyladenosine in Vasoactive microRNAs during Hypoxia; A Novel Role for METTL4. Int J Mol Sci 2022, 23.

38. Lichinchi G, Gao S, Saletore Y, Gonzalez GM, Bansal V, Wang Y, Mason CE, Rana TM: Dynamics of the human and viral m(6)A RNA methylomes during HIV-1 infection of T cells. Nat Microbiol 2016, 1:16011.

39. Lichinchi G, Rana TM: Profiling of N(6)-Methyladenosine in Zika Virus RNA and Host Cellular mRNA. Methods Mol Biol 2019, 1870:209–218.

40. Lichinchi G, Zhao BS, Wu Y, Lu Z, Qin Y, He C, Rana TM: Dynamics of Human and Viral RNA Methylation during Zika Virus Infection. Cell Host Microbe 2016, 20:666–673.

41. Platanias LC: Mechanisms of type-I- and type-II-interferon-mediated signalling. Nat Rev Immunol 2005, 5:375–386.

