## Supplementary figures and images for "Cannabinoids shift the basal ganglia microRNA m^6^A methylation profile towards an anti-inflammatory phenotype in SIV-infected rhesus macaques"

### Supplemental Figures 1 to 3

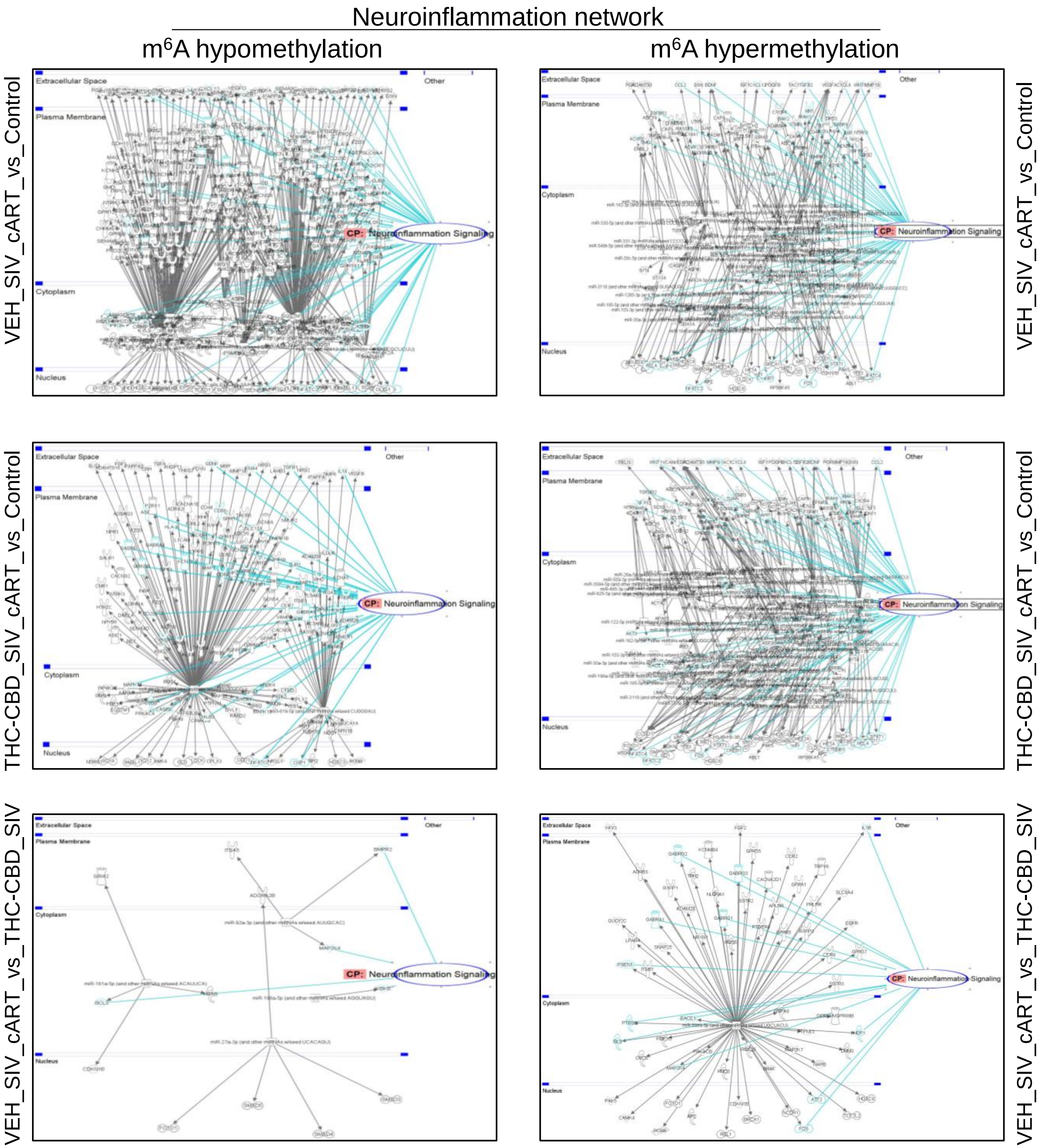

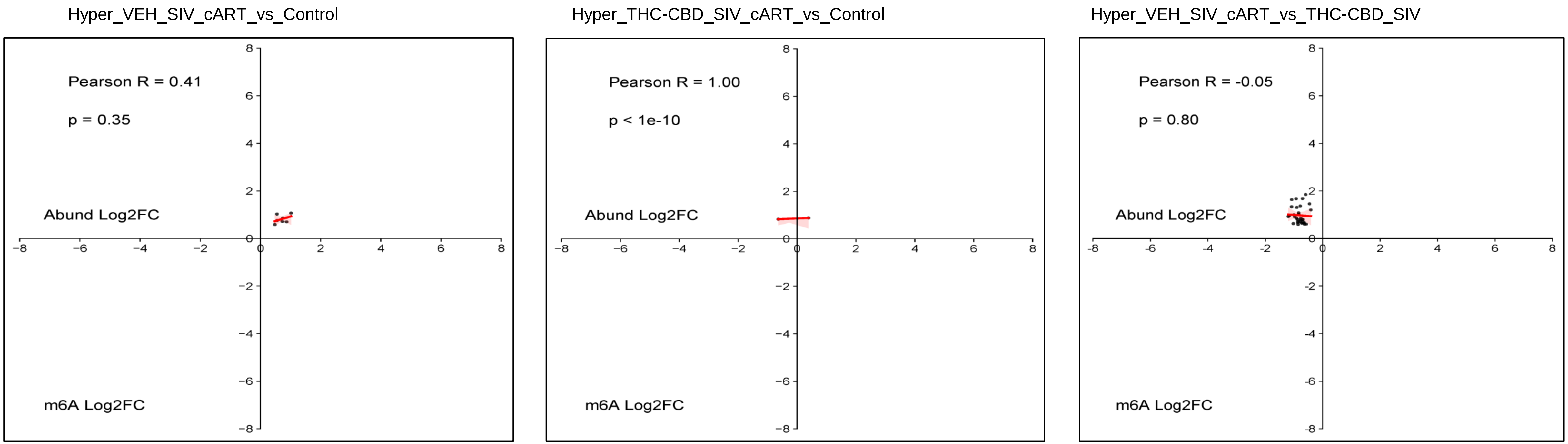

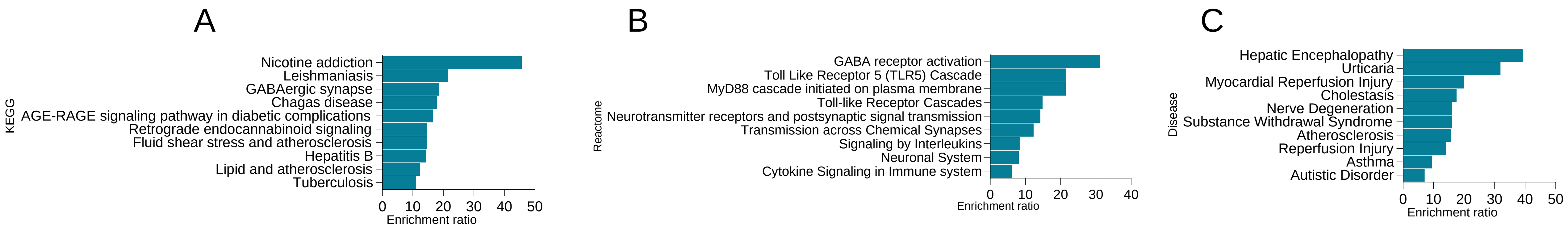
